# Agent-based modelling and time series inference of filamentous yeast colonies

**DOI:** 10.1101/2025.09.15.676253

**Authors:** Kai Li, Alexander K. Y. Tam, Jennifer M. Gardner, Joanna F. Sundstrom, Vladimir Jiranek, J. Edward F. Green, Benjamin J. Binder, Andrew J. Black

## Abstract

The baker’s yeast *Saccharomyces cerevisiae* can form invasive filamentous colonies with non-uniform spatiotemporal patterns. We aimed to better understand how individual cellular actions give rise to colony-scale patterns. We used an off-lattice agent-based model to simulate colony growth, and used a neural likelihood estimator (NLE) to infer parameters for experimental photographs. Li et al. [1] used approximate Bayesian computation (ABC) to infer parameters using coarse summary statistics obtained from a single time point averaged across experimental replicates. The NLE overcomes the computational expense of ABC, allowing us to infer the parameters of individual colonies from a full time series of experimental photographs. To demonstrate the capabilities of our model and inference technique, we tested extensively on synthetic data and then predicted yeast growth under three different experimental conditions. As before, the proportion of total colony growth above which pseudohyphal growth is permitted was a key parameter that contributed to colony morphology. Since our NLE-based approach incorporates time series data, it yielded better parameter estimates and more accurate predictions compared to our ABC-based method. This updated approach improved understanding of how the probability that a stated cell produces a pseudohyphal cell influences colony morphology. In this way, the model also has the potential to generate hypotheses, which can be tested through biological experiments to increase the understanding of the basis for different growth patterns in yeast.

## 1 Introduction

Yeasts are among the most versatile microorganisms studied in biology. Due to their low cost and ease of culturing, yeasts are often used as a model organism for other eukaryotic cells in scientific research [2]. With over 1,200 known species, yeasts also play a vital role in human life. For example, the yeast *Candida albicans* is an opportunistic pathogen that can cause serious infections in humans [3]. These infections are underpinned by yeast’s ability to invade surrounding tissue. Yeasts also have positive impacts. For example, the baker’s yeast *Saccharomyces cerevisiae* is used in the production of baked goods and alcoholic beverages [4]. Other positive applications of yeasts include in the biotechnology industry for biofuels [5], in the health industry for drug discovery [6] and in advancing genome research [7]. There is also ongoing research in optimising and developing new strains for producing specialised wines [8], for example the AWRI 796 and Simi White strains involved with this work. Hence, there is great motivation in studying yeast.

Filamentous *S. cerevisiae* colonies consist of two cell types [9], which we term sated and pseudohyphal. Sated cells are ellipsoids with average major axis lengths of 4.2 µm and minor axis lengths of 3.0 µm. Pseudohyphal cells are also ellipsoidal, but are elongated compared to sated cells, with axis lengths of 6.7 µm and 1.9 µm [1, 9]. Sated cells bud from sites located near both ends of the major axis, whereas pseudohyphal cells bud from the distal pole only [10]. Budding is an asexual process driven by nutrients. In filamentous colony experiments, nutrient diffusion is much faster than cell proliferation [11]. Consequently, the nutrient concentration is effectively uniform in space, but depletes over time due to consumption by cells. Yeast cells are non-motile, and since nutrients are spatially uniform, they do not undergo directed growth [11], as the roughly symmetrical colonies of Figure 1.1 indicate. During early growth, when nutrients are abundant, *S. cerevisiae* colonies exhibit spatially uniform Eden-like circular growth. This circular growth consists of sated cells only, and is shown in Figure 1.1(a)–(b). However, when nutrients deplete the colony produces elongated pseudohyphae (Figure 1.1(c)–(h)) in an attempt to access more nutrients. Pseudohyphal growth also occurs underneath the colonies, where cells invade their substratum [12]. This invasive phenomenon is important clinically, but in this work we restrict ourselves to two-dimensional surface growth.

**Figure 1.1.**
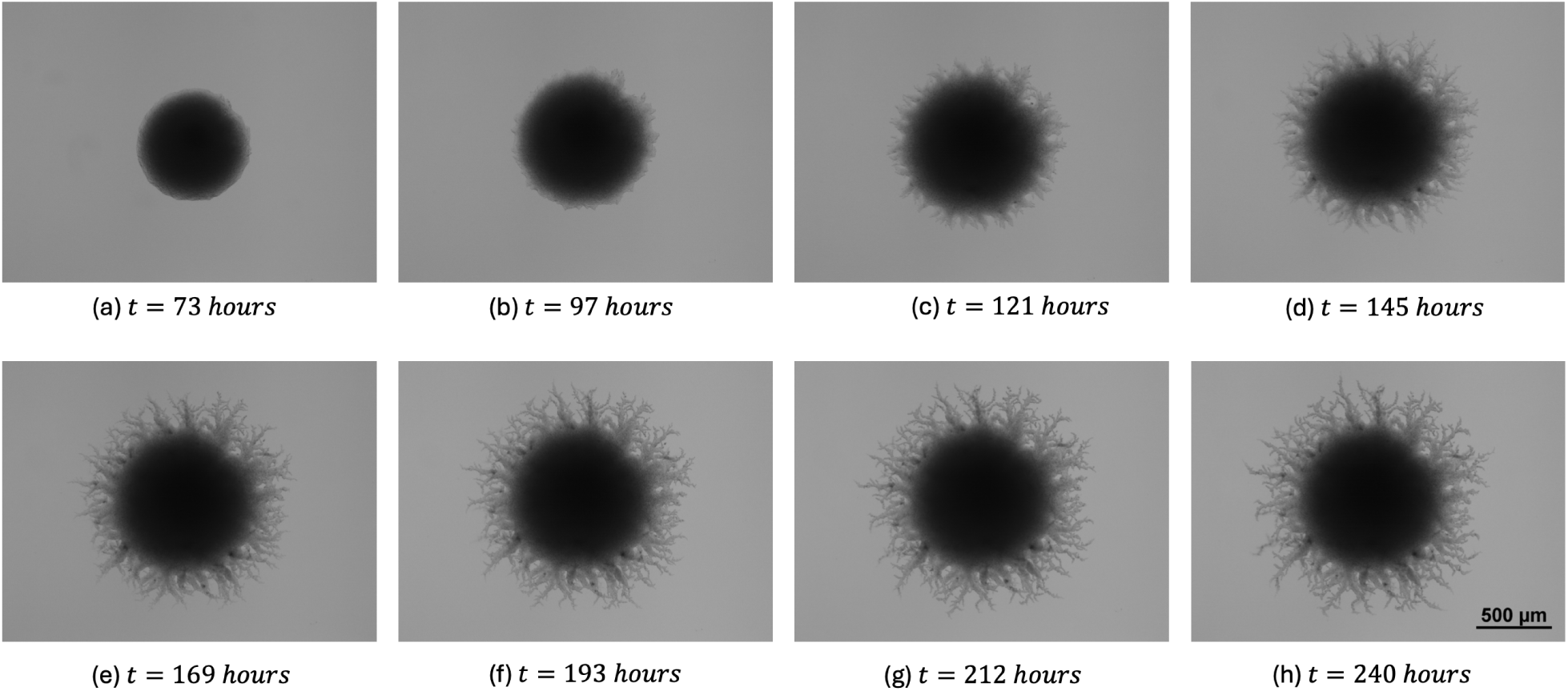
Time series images of a growing yeast colony of AWRI 796 50 µm showing the transition from Eden-like circular growth to filamentous growth.

In this paper, we use agent-based modelling to simulate colony-level morphologies arising under different experimental conditions. Agent-based models have been used extensively in biology to study cancer [13–15], tissue growth [16–18], molecular biology [19–21], as well as yeast [11, 22, 23]. Li et al. [1] used an off-lattice agent-based model, where the agent represents a cell, to capture filamentous yeast growth. Li et al. [1] confirmed the important role of a cell phenotype switch from sated to pseudohyphal in colony morphology. In our model, this switch occurs when the colony attains a threshold size, which promotes filamentation by depleting surrounding nutrients, and requiring more resources to sustain the larger colony. We return to the model of Li et al. [1] in this paper, with the key extension of using time series data for individual colonies to parameterise the model. This time series data consist of a series of 8 photographs of individual colonies at different stages of development. Incorporating the additional data extends the work of Li et al. [1], who used data from a single time point at the end of the experiment, and summary statistics averaged over experimental replicates. Analysing the time series data reveals that the probability that a stated cell produces a pseudohyphal cell is a key determinant of colony morphology, which was not fully appreciated using data from only a single time slice.

Since approximate Bayesian computation (ABC) [24] is expensive and scales poorly to large data sets, our time series data requires a more computationally efficient inference method. Hence, in this work we perform inference using neural likelihood estimation (NLE) [25–27]. NLE is an inference method that uses a conditional density approximation parameterized by a neural network to approximate the likelihood function. A key advantage of neural likelihood methods is that they require significantly fewer simulations to train the model compared to doing inference with ABC [25], and can more easily exploit parallel computing resources. NLE is most useful where the likelihood is intractable and model simulation is expensive, and has found applications in economics [28], cosmology [26] and neuroscience [29]. Since neural methods drastically speed up parameter inference compared to ABC, in this paper we explore the variability of parameter values within an entire set of 14 individual experiments for a given yeast strain and initial nutrient level. This process required less than a day of computational time, whereas with the ABC method this would have taken many months due to the high computational cost of simulation, image processing, and parameter inference.

## 2 Materials and Methods

Our approach combines yeast growth experiments, mathematical modelling, image processing, and parameter inference. Subsequent subsections outline our methods for each of these steps.

### 2.1 Yeast Growth Experiments

We use the same experimental set-up as described by Li et al. [1]. The experiments are performed using the AWRI 796 strain at two initial ammonium sulfate nutrient concentrations (AWRI 796 50 µm and 500 µm), and the Simi White strain at 50 µm. These are two commercially available strains of *S. cerevisiae* used in wine production. While colonies are growing, we take a photograph of each daily, from Day 3 (73 hours) until Day 10 (233 hours). Example photographs from one replicate are shown in Figure 1.1. We have photographs for 14 individual colonies for each set of experimental conditions. When referring to a set of experiments, we name them according to yeast strain and nutrient level. For example, AWRI 796 50 µm refers to experiments of the yeast strain AWRI 796, with an initial nutrient concentration of 50 µm.

### 2.2 Mathematical Model

We use the two-dimensional off-lattice agent-based model for a filamentous yeast colony developed by Li et al. [1]. The model contains two cell types: sated and pseudohyphal, and assumes that all cells of a given type are ellipses with the same dimensions: axis lengths of 4.2 µm and 3.0 µm for sated cells, and 6.7 µm and 1.9 µm for pseudohyphal cells. In practice, we approximate both cell types as dodecagons. We simulate colony growth one cell division event at a time starting from a single cell, until the number of cells in the colony, *n,* reaches a maximum value prescribed in advance, *n*_max_. This value *n*_max_ is chosen to approximate the number of cells in the experimental or synthetic colony that we seek to simulate. Since we do not model nutrients explicitly, this assumption enables us to neglect time, removing the need to have unknown proliferation rates as additional parameters to be inferred. Sated cells proliferate from one of 4 possible budding sites located at both ends of the cell, whereas pseudohyphal cells can only bud from one of 2 sites located at prescribed angles from the distal pole.

A schematic of the two cell types and all possible cell proliferation events in the model is shown in Figure 2.1. Sated cells are indicated in green, and pseudohyphal cells are indicated in purple. Our time-free model contains five key parameters that influence filamentous yeast growth, each taking values on the unit interval [0, 1]. In vector form, we write the parameters as ***θ*** = (*n*^∗^*, p_a_, p_sp_, p_ps_, γ*), and describe their meanings below.

**Figure 2.1:**
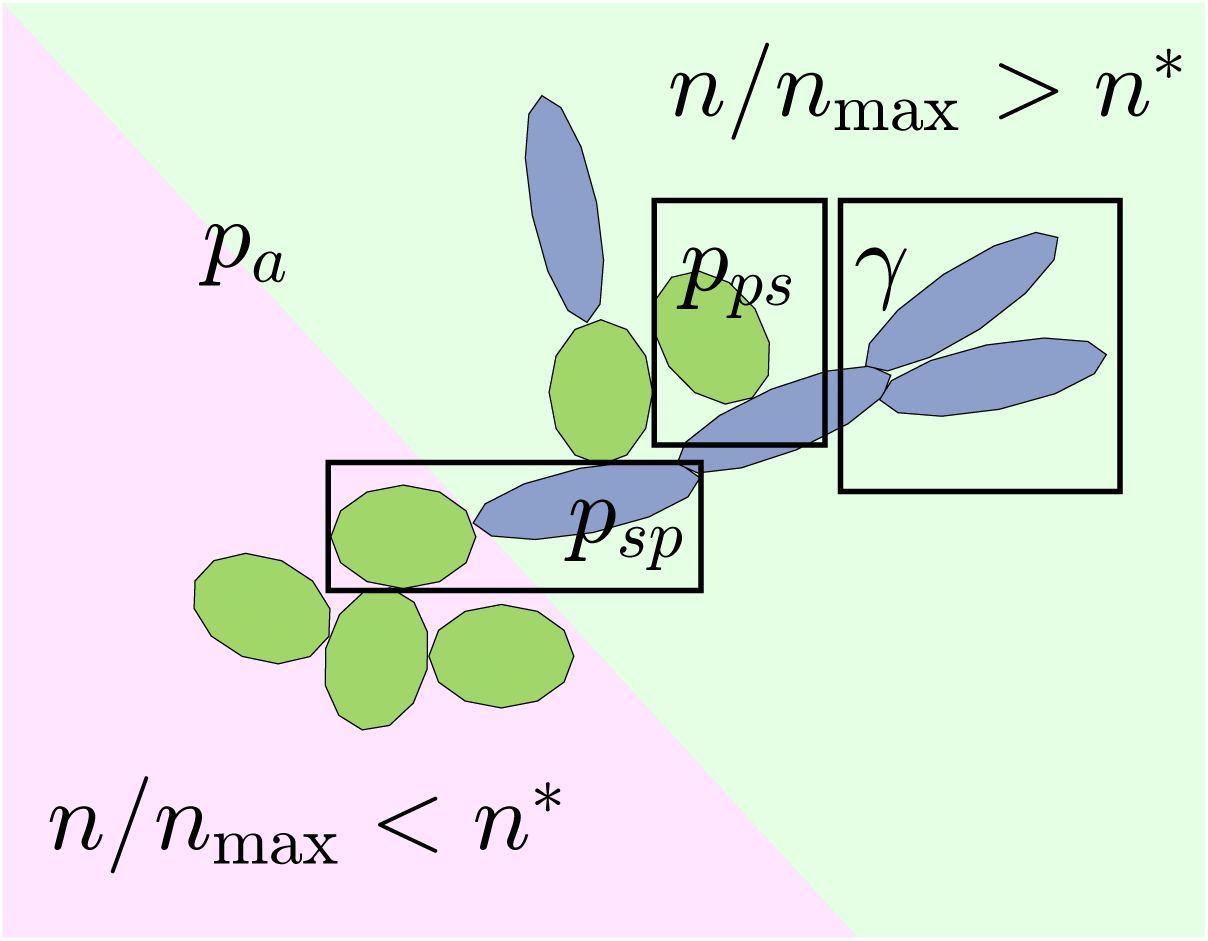
Budding patterns in the agent-based model. All cells are sated (green dodecagons) until the threshold *n > n*^∗^*n*_max_ is attained. After this threshold, the first pseudohyphal cell (blue dodecagons) can be produced. Then, more sated and pseudohyphal cells are produced via a set of proliferation events.

For each individual colony simulation, we assume a threshold number of cells, *n*^∗^*n*_max_, below which all cells are sated and no pseudohyphal growth occurs. This sated-only growth is shown in the pink region of Figure 2.1. If the current number of cells in the simulation *n > n*^∗^*n*_max_, pseudohyphal growth becomes possible. We then implement a series of steps to determine the next cell to proliferate, and the type of daughter cell produced. If one or more pseudohyphal cells are present, then the next cell to proliferate will be a sated cell with probability *p_a_*, or a pseudohyphal cell with probability 1 − *p_a_*. Once a cell type is chosen, the proliferating cell is chosen at random from all cells of that type. If a sated cell is chosen, it will produce a pseudohyphal daughter cell with probability *p_sp_*, or a sated cell with probability 1 − *p_sp_*. If a pseudohyphal cell is chosen and does not have any daughter cells, then a pseudohyphal daughter cell is produced from the distal pole. However, if the pseudohyphal cell chosen to proliferate already has an existing pseudohyphal daughter cell, its behaviour is controlled by two parameters, *γ* and *p_ps_*. If the selected pseudohyphal cell has a pseudohyphal daughter, we abort proliferation with probability *γ* and instead select a new pseudohyphal mother cell with no pseudohyphal daughter to proliferate. Larger values of the parameter *γ* limit forking, which is the scenario where multiple pseudohyphal branches emanate from the same pseudohyphal mother. Proliferation proceeds with probability 1 − *γ*, and the daughter cell is sated with probability *p_ps_*, and another pseudohyphal cell with probability 1 − *p_ps_*. If at any stage in the simulation the cell chosen to proliferate has no budding site available or if the centre of the daughter cell would lie within the boundary of another cell, proliferation is aborted, and the procedure begins again from the start.

### 2.3 Image Processing and Summary Statistics

Image processing and quantification enabled us to compare experimental and simulation results. The first step was to convert grayscale photographs and simulation plots to binary images. We obtained the binary images using the Tool for Analysis of the Morphology of Microbial Colonies (TAMMiCol) [30], which uses automatic thresholding and interpolation to obtain binary images. We saved simulation images at the same pixel resolution as experimental photographs. Computing summary statistics for the spatial patterns in images enabled us to compare simulated and experimental images. We compute six summary statistics per image: colony area, maximum radius, mean radius, branch length, filamentous area, and compactness. These summary statistics quantify the size, extent of filamentous growth, and spatial pattern of the filaments.

The colony area (*I_A_*) is the total number of occupied pixels in the binary image, which is readily obtained in Matlab. To compute the maximum and mean radii, we first used Matlab to locate the colony centroid. The mean radius (*I_R_*_mean_) is the mean distance between the centroid and any pixel located on the colony perimeter, and the maximum radius (*I_R_*_max_) is the maximum distance from the centroid to an occupied pixel. The mean radius indicates the amount of the colony occupied by the central circular region of Eden-like growth. We achieved better results using the mean radius (see Figure 2.2(b)) to quantify the size of the circular region than the complete spatial randomness [31] and minimum [1] radii, which we have used in our previous work.

**Figure 2.2.**
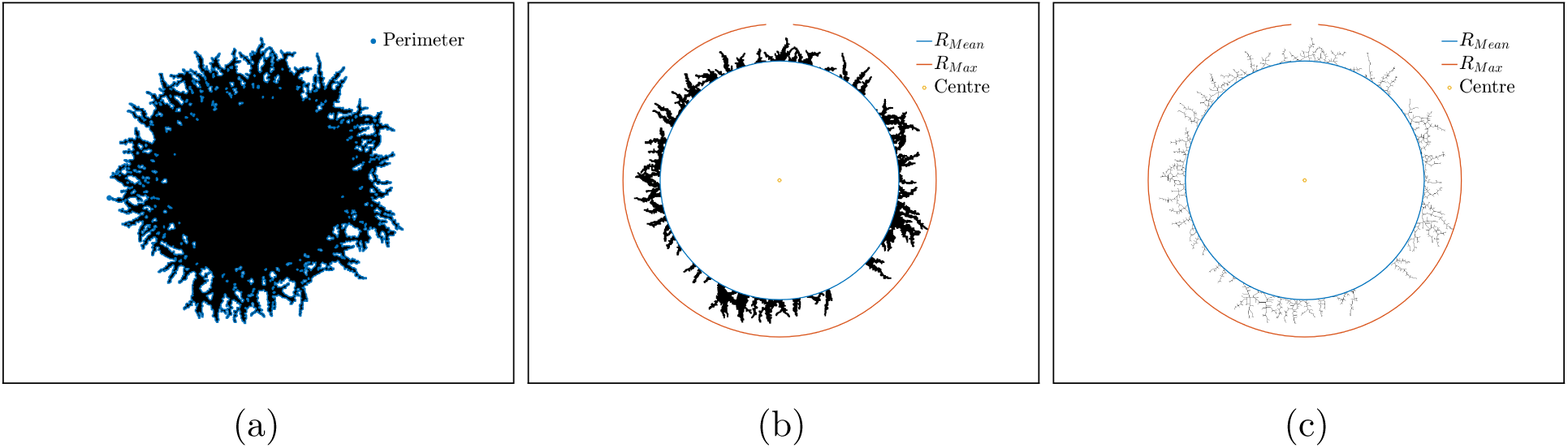
Visualisation of summary statistics for a binary image from a simulated colony. (a) Binary image with perimeter overlayed. (b) Processed binary image indicating the colony centroid, maximum radius, mean radius, and filamentous area. (c) Skeletonized binary image of the filamentous region, which was used to compute the branch length.

The filamentous area (*I_A_*) is the number of occupied pixels in the annular region between the mean radius and the maximum radius. Filamentous area quantifies the extent of filamentation but also accounts for the density of filaments. To obtain the branch length (*I_B_*), we summed the length of branches of a skeletonised experimental image, an example of which is shown in Figure 2.2c. Matlab’s bwmorph() function provides a count of the number of branches in a skeletonised image of the filamentous region. Due to the computational cost of skeletonisation, we only counted a quarter of the colony’s branches. Finally, compactness (*I_C_*) is the colony’s perimeter-to-area ratio, *P/A,* normalised by the perimeter-to-area ratio of a circle with area *A*. Compactness therefore quantifies the irregularity of the colony shape, such that a perfect circle has the smallest possible value *I_C_* = 1, and deviations from the circle result in *I_C_ >* 1. Using the colony perimeter (see Figure 2.2(a)) and area obtained from Matlab, we compute the compactness using *I_C_* = *P* ^2^*/*(4*πA*).

To show how these summary statistics vary throughout experiments, we plotted each summary statistic from Day 3 (∼ 73 hours) until Day 10 (∼ 233 hours). An example of these six summary statistics evaluated on time series data of AWRI 796 50 µm colonies is given in Figure 2.3. Each semi-transparent line represents an experimental colony, with 14 colonies in total. We also overlay these with box plots at each recorded time point to show the spread of the data, with the red crosses indicating outliers.

**Figure 2.3:**
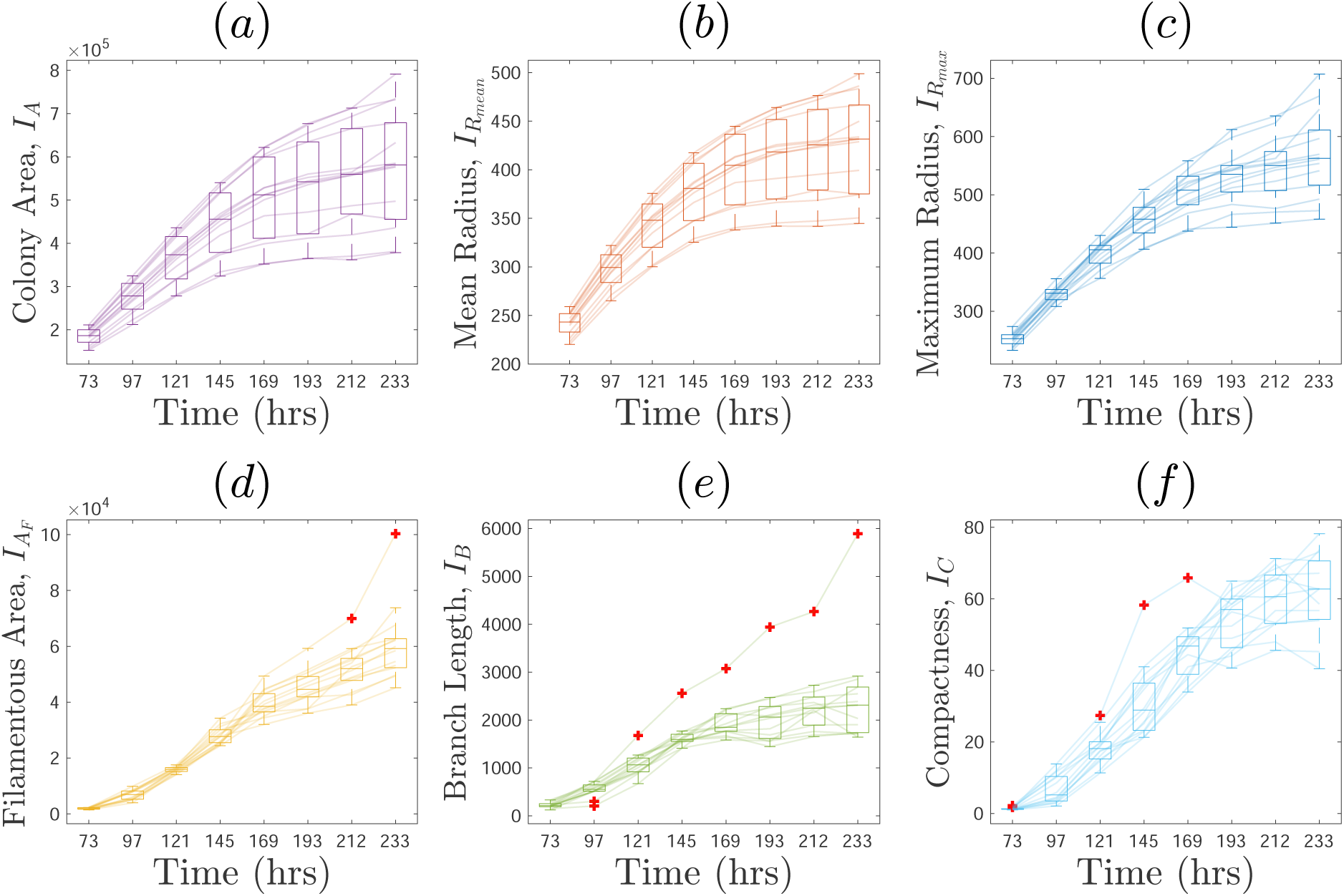
Summary statistics used to quantify AWRI 796 50 µm colonies over time. (a) Colony area, *I_A_.* (b) Mean radius, *I_R_*_mean_ . (c) Maximum radius, *I_R_*_max_ . (d) Filamentous area, *I_A_* . (e) Branch length, *I_B_*. (f) Compactness, *I_C_*.

### 2.4 Parameter inference

Given observed data ***t***^′^ from a series of experimental images, our goal was to estimate the posterior distribution of the parameters *p*(***θ***|***t***^′^) ∝ *p*(***t***^′^|***θ***)*p*(***θ***), where *p*(***θ***) is the prior distribution and *p*(***t***^′^|***θ***) is the likelihood. The simulation described in the previous section gives a way of sampling from the distribution *p*(***t***|***θ***), but actually evaluating it is intractable. NLE instead takes advantage of this ability to simulate the model to learn an approximation to the likelihood function. We can then sample from the posterior using standard Markov chain Monte Carlo (MCMC) methods, by replacing the true likelihood with the approximation.

Starting from a set of samples {***θ****_i_, **t**_i_*}*_i_*_=1:*N*_ that are obtained by ***θ****_i_* ∼ ̃*p*(***θ***) and ***t****_i_* ∼ *p*(***t***|***θ_i_***), we trained a conditional neural density estimator *q_w_*(***t***|***θ***), where *w* are the weights of a neural network. The proposal distribution ̃*p*(***θ***) controls where the data is generated, and therefore how accurate the approximation is. This introduces some circularity to the problem as we would like ̃*p*(***θ***) to concentrate in areas of high posterior density, but we do not know this *a priori*. To address the issue of circularity, current state-of-the-art methods take a sequential approach to generating training data and refining the proposal [25]. Our testing revealed that this sequential approach was unnecessary for our problem, and good results could be obtained from a single round of training on a sufficiently large dataset. This is probably due to the simple shape and limited support of the posterior distributions in our problem.

The conditional neural density estimator that we use in this work is a Gaussian mixture density network (MDN). The conditional density takes the form

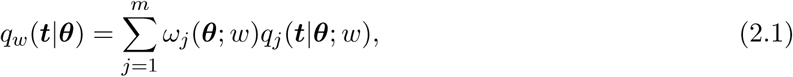

where *ω_j_*(***θ***; *w*) are the mixture weights such that ^L^*_j_ ω_j_* = 1 and *q_j_*(***t***|***θ***; *w*) are multivariate normal densities *q_j_*(***t***|***θ***; *w*) = N (***µ****_j_*(***θ***; *w*), Σ*_j_*(***θ***; *w*)). The weights, means and covariances are all functions of the parameters ***θ***, which are in turn determined by the weights of the neural network *w* [32]. We implement the MDN using the Julia package MixtureDensityNetwork.jl. Below are detailed steps for how we trained and sampled from the models.

The parameter vector ***θ*** has five elements, and each parameter takes values on the unit interval. We generate the training dataset from independent uniform distributions, *i.e*. ̃*p*(***θ***) = 1, ***θ*** ∈ [0, 1]^5^. The domain of the MDN is R^5^, so we apply a logit transform to the samples such that the transformed parameters take values on an unrestricted domain when training the model. Note that the Metropolis-Hastings (MH) sampling discussed later is also performed on this unrestricted space, and then samples are transformed back to [0, 1]^5^ to visualise the final posteriors. This requires the addition of Jacobian terms in the calculation of the MH acceptance probabilities that are detailed in Appendix A.1.

We define the components of the observed data t as

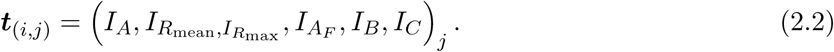

where *i* indexes the six different summary statistics [*I_A_, I_R_*_mean_*_,IR_*_max_ *, I_AF_, I_B_, I_C_*] and *j* = 1, · · · *, T_F_* is over the number of time points. Typically, we set *T_F_*= 8 as there is a maximum of eight time course images per colony. For example, to interpret the component of the vector ***t***, we take ***t***_(1,8)_, which is the area of the colony at time point eight. As the summary statistics vary by different orders of magnitude, as shown in Figure 2.3, we normalised within each summary statistic by dividing by the maximum value of each time series summary statistic from the simulations of the prior. This ensures all summary statistics are close to the unit interval while preserving the trend of the time series data. Note that for a set of simulations, we simulate up to a target cell number which is prescribed in advance based on the experimental colony area. Hence, the summary statistics *I_A_* will be approximately the same for all simulations and the target colony.

The architecture of the neural networks and the number of mixtures were determined by splitting the data into a 90/10 training/test set and performing a number of experiments monitoring the log likelihood. From this we found that training the MDN using 600 epochs, a batch size of 16, two hidden layers of depth 64 each, and a mixture of six yielded good results across all experimental and synthetic conditions, so this architecture was fixed. Throughout we use the Adam optimiser with a learning rate of 1 × 10^−3^. The MDN model was trained on 1000 simulated data points. This number of simulations was chosen because they could be completed on the available HPC resources in less than 24 hours. This lent itself to a convenient workflow for running and analysing results without requiring a sequential method, which although probably a more efficient use of resources requires more care in tuning [25].

After learning the likelihood function, we sample from the posterior distribution using a basic MH algorithm. The proposals were independent Normal distributions with a standard deviation of 1. We took 200,000 samples to estimate the posterior distribution and remove the first 10,000 samples of burn-in. Thinning was also applied to the chains by taking every tenth parameter value to reduce autocorrelation. This part of the procedure is very fast, so it did not require any substantial tuning. As discussed above, the final step was to apply an inverse logit transform to the samples to return them to the unit interval.

## 3 Results

### 3.1 Synthetic data set

To test how well our inference algorithm could uncover the true parameter values, we simulated synthetic datasets and performed inference to recover the model parameters. We show one example here in the main text and another two in the Appendix. Instead of initialising the synthetic test colony from random parameters, we used the parameter values obtained by Li et al. [1] for AWRI 796 50 µm. We did this because non-physical colony morphologies will result from many combinations of random parameters. Since our summary statistics were designed with the specific aim of capturing the morphology of filamentous yeast colonies, they work best when the colonies are similar to the experimental colonies. Therefore, non-physical colonies are not suitable for the analysis. For the example shown here, the parameters used were *n*^∗^ = 0.67 *p_ps_*= 0.58 *p_sp_* = 0.25, *γ* = 0.12, *p_a_* = 0.14. We simulate the synthetic colony up to the same area as observed in those experiments for AWRI 796 50 µm sample one (see Figure 1.1) and we use all six summary statistics at all eight time points, providing 48 summary statistics per colony.

A pair plot of the posterior distribution is shown in Figure 3.1, where the ground truth parameters are shown with blue lines. The proportion of total colony growth above which we permit pseudohyphal growth, *n*^∗^, could be recovered with a small variance. This parameter has an important impact on colony morphology, as it determines when filamentation can begin. For other parameters, such as *p_ps_* and *p_sp_*, the true values lie in the tails of the distribution. The forking parameter *γ* and the probability of biasing sated cell proliferation, *p_a_*, lie near the mode of the posterior. These parameters could be recovered with some uncertainty due to the large variance in the distributions.

**Figure 3.1:**
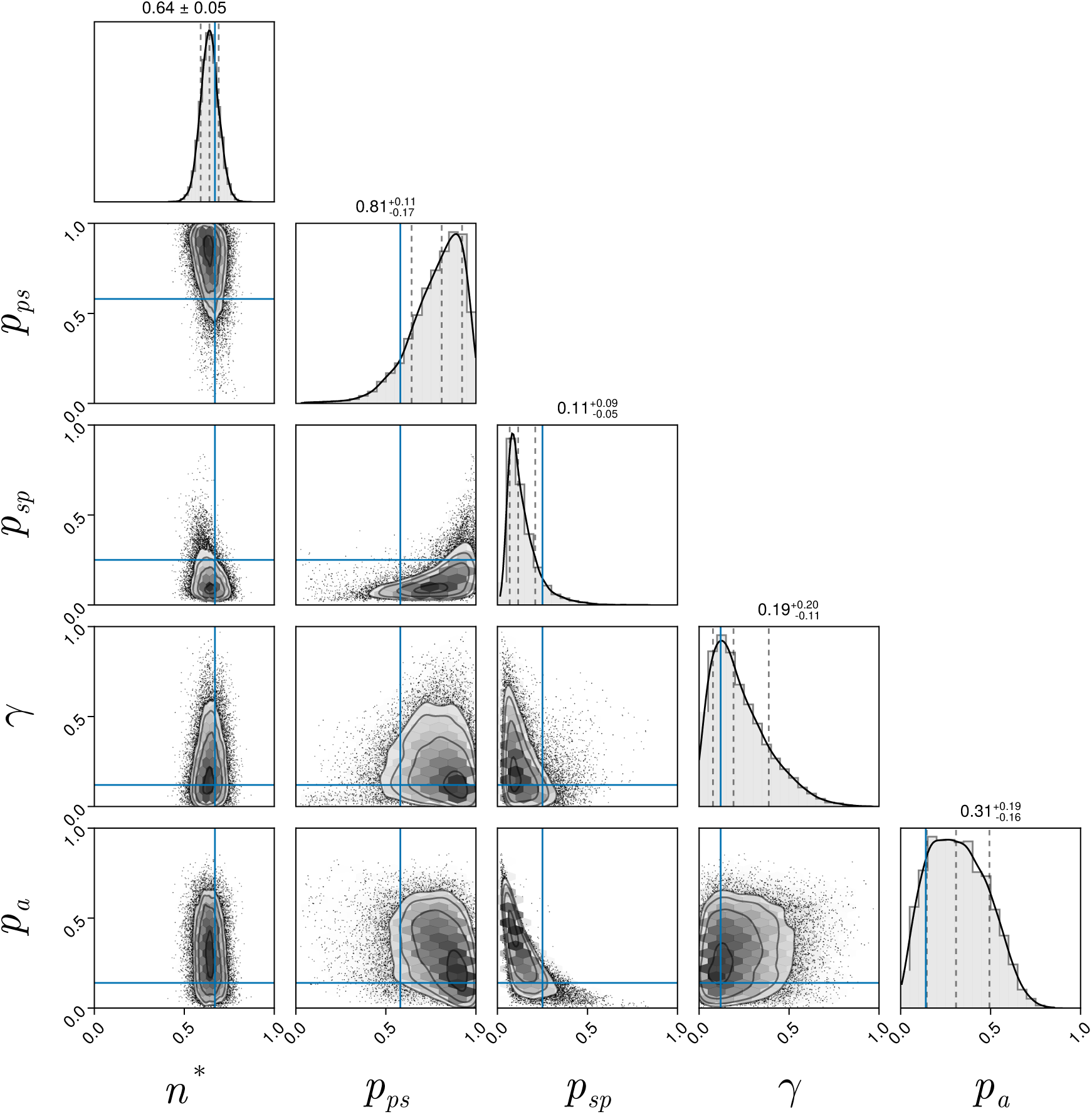
Pair plots of posterior distribution for the synthetic data. The blue line represents the ground truth parameter values. *n*^∗^ = 0.67 *p_ps_* = 0.58 *p_sp_* = 0.25, *γ* = 0.12, *p_a_* = 0.14.

**Figure 3.2:**
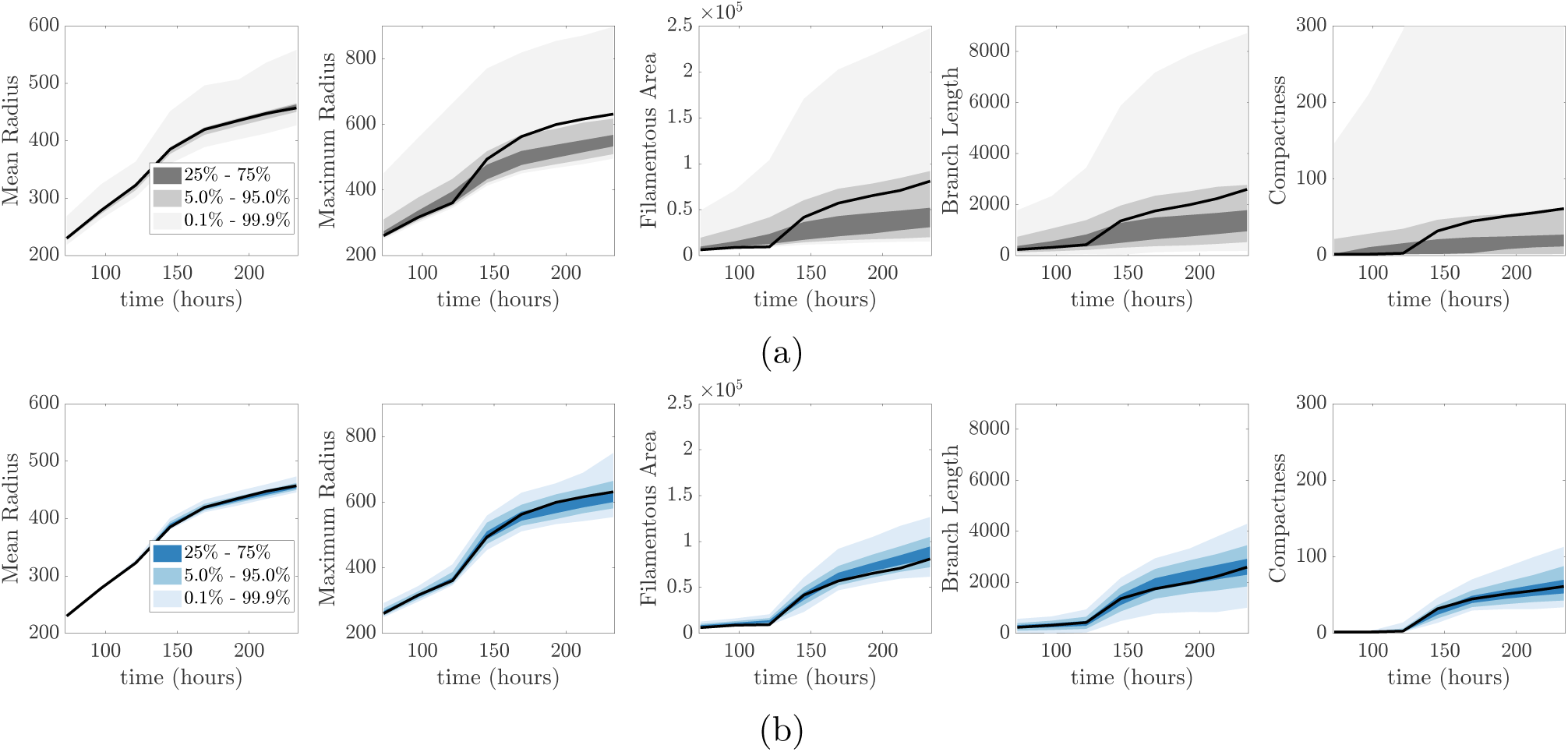
Ribbon plots showing the prior (a) and posterior (b) predictive distribution for the synthetic dataset. The black lines shows the realisation used for the inference.

**Figure 3.3:**
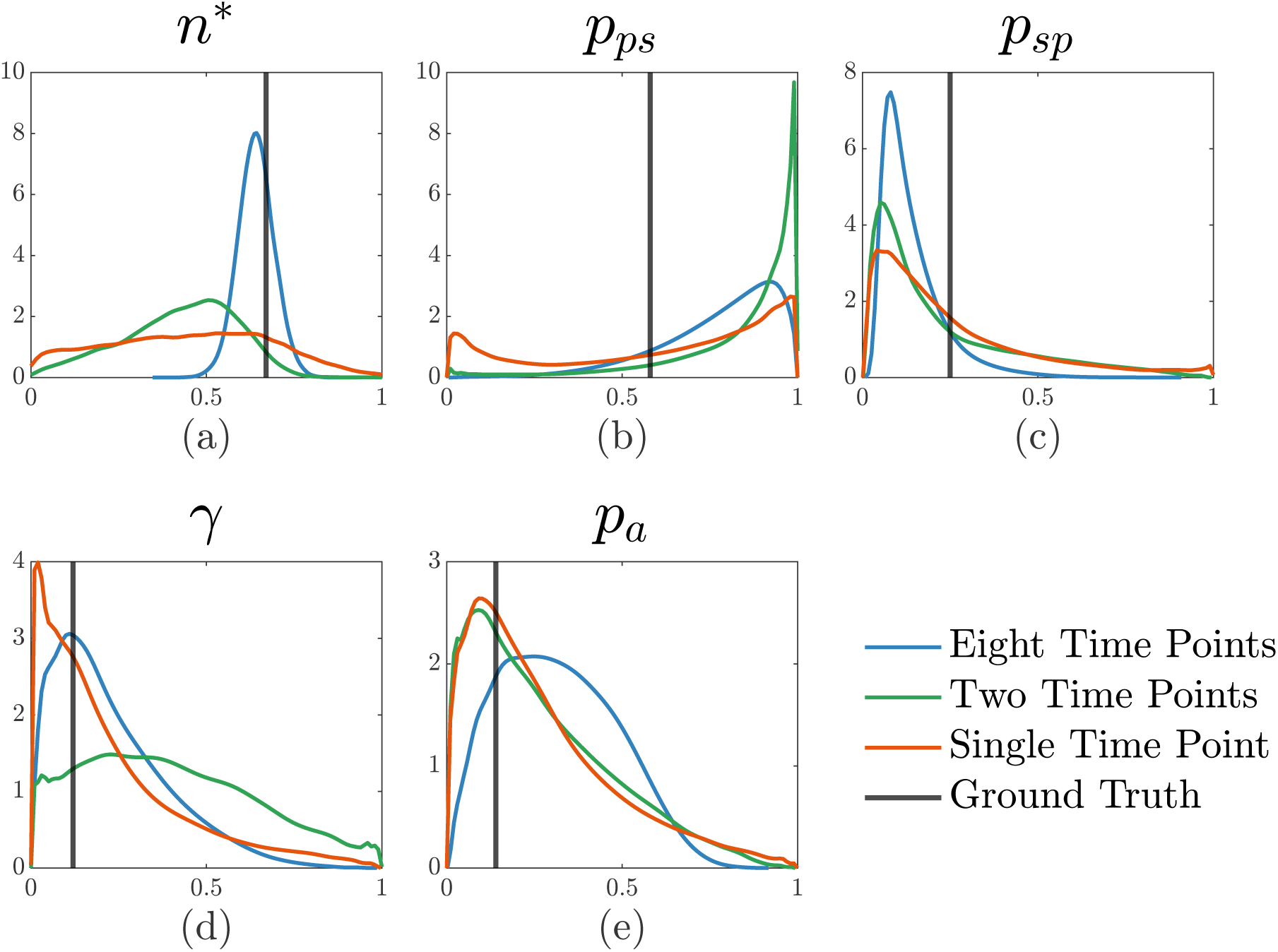
Marginal posterior distributions from performing inference using data from an increasing number of time points.

**Figure 3.4:**
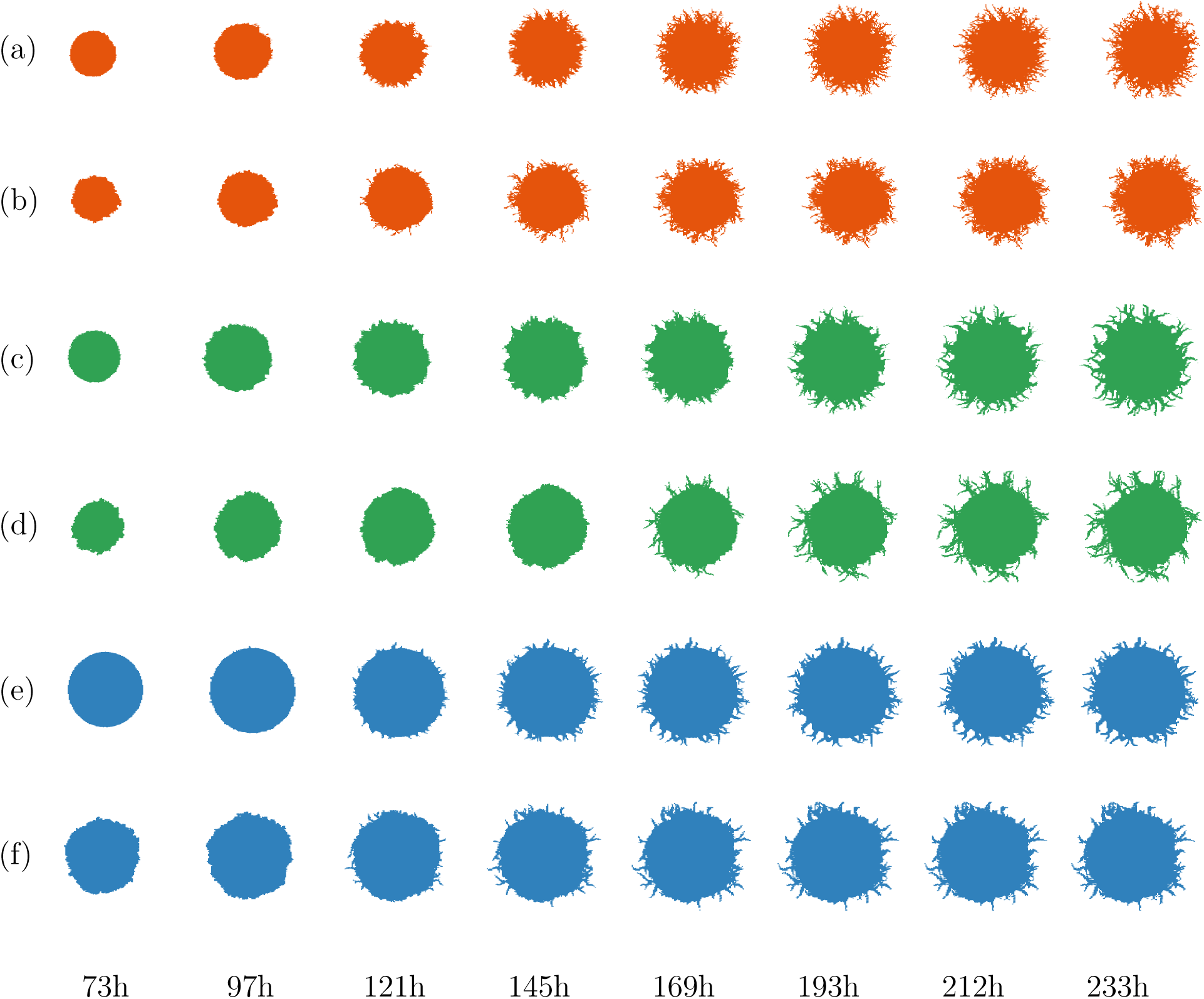
Colony simulation comparison between experiment and posterior prediction. (a) Experiment of AWRI 796 50 µm Sample 1 filamentous yeast. (c) Experiment of Simi White Sample 7 50 µm filamentous yeast. (e) Experiment of AWRI 796 500 µm Sample 9 filamentous yeast. (b,d,f) Respective simulations with parameter values sampled from the posterior distribution in Figure 3.5 with colours distinguishing between different experiments.

To check the performance of the inference we also simulated colonies by sampling parameters from the prior and posterior distributions and created ribbon plots of the summary statistics over time. As expected, the synthetic datasets lie within both the prior (Figure 3.2a) and posterior (Figure 3.2b) predictive intervals as the realisation is drawn from the prior distribution. After approximately 125 hours, filamentous area, branch length, and compactness increased sharply. These increases probably occur because *n > n*^∗^*n*_max_ after this time, initiating filamentous growth. Moreover, the posterior prediction distribution is centred around the true summary statistics and has much less variability than the prior predictive distribution. This demonstrates that the inference is performing as expected, even though some of the parameters are only weakly inferred.

To further validate the parameter inference, we randomly sampled 10 sets of parameters from the posterior distribution and plotted images of the simulated colonies through time to provide a visual comparison with the colony used for the inference. The 10 synthetic colonies shared very similar morphologies to the original data (see Appendix Figure A.5). The yeast colony model is stochastic so there is some natural variability in the summary statistics for a fixed set of parameters. This can be seen in Figure A.2, which shows ten realisations grown to the same area for the same fixed parameter values. We investigated how this variability affects the inferred parameters by rerunning the inference on these trajectories, and the posteriors were broadly similar.

We further tested our inference on two more synthetic data sets with different parameter values to test the robustness of our inference algorithm. For these data sets we used smaller values of *n*^∗^ (0.28 and 0.39) as this will induce filamentation earlier compared to the first synthetic data set; other parameters were also varied. The posterior pair plots for both are shown in Figure A.1a. We also present their respective predictive distribution in Figure A.3. Again, we could retrieve *n*^∗^ to a high degree of accuracy, while the other four parameters can also be somewhat inferred with the ground truth near the mode of the posterior distributions. For both *n*^∗^ = 0.28 and *n*^∗^ = 0.39, the ground truth summary statistic values (solid black line) lie within the posterior predictive intervals. Moreover, due to the lower variance of posterior distributions compared to the example shown above, we see much narrower posterior predictive intervals compared to the prior predictive intervals. For a smaller *n*^∗^, the other parameters are more identifiable. This is because a smaller value of *n*^∗^ triggers earlier filamentation and since many of the summary statistics quantify filamentation, this makes the other parameters more identifiable.

#### 3.1.1 Increasing the number of summary statistics improves parameter inference

We next investigated how the addition of time-series data improved the parameter inference over just using a single image at the end of the growth period as in Li et al. [1]. We reran the inference on the synthetic data using data from just the last time point (using the notation defined in Section 2.4 this is **t**_(*i,*8)_, for all *i* = 1*, . . .,* 6), two time points (**t**_(*i,*4)_ and **t**_(*i,*8)_), and all eight (**t**_(*i,j*)_, for all *j* = 1*, . . .,* 8). The marginal posteriors for this experiment are shown in Figure 3.3.

The most clear result is that the posterior for *n*^∗^ contracts sharply with more data becoming much more informative. Finer resolution data allows the model to narrow down when filamentation begins, allowing for a much better estimate of *n*^∗^. Similarly, *γ* is more identifiable as the number of summary statistics increases. This is evident by a decrease in the variance and the ground truth (blue vertical line) being situated around the mode of the distributions. For *p_sp_*, there was little change in the mode of the distribution, with the true value still lying in the tail of the distribution. Moreover, *p_a_* increased in uncertainty as the number of summary statistics increased. This could be because learning the higher-dimensional distribution is harder, or could be just due to the stochasticity in the time series. The parameter *p_ps_* had large variability across all three types of summary statistics, suggesting this parameter is unidentifiable, which is consistent with the results of Li et al. [1].

Figure A.7 shows the posterior prediction ribbon plots for these experiments. These clearly show that more data improves the accuracy of the posterior predictions across all time points of the simulation. The effect of a tighter posterior distribution on *n*^∗^ can be seen visually in Figure A.8. More data can accurately constrain the time at which filamentation begins producing better agreement near the start of the time series. Hence, we use all eight images when running the inference.

### 3.2 Experimental modelling and inference

Having tested our inference algorithm on synthetic datasets, we turn to the experimental colonies for the three experimental conditions. This allowed us to gain further insight into the biological mechanism contributing to the colony morphology development of *S. cerevisiae* at different stages in time. Overall, the model performs well in capturing the colony morphology. Figure 3.4 compares the experimental images with simulated colonies where parameters are randomly sampled from the corresponding posteriors. We also present an additional ten simulations in the Appendix A.4. Visually, the simulated colonies match very closely to the experimental colonies at all time points, indicating that our model predictions effectively capture the colony morphology.

We plot the prior and posterior predictive distributions as ribbon plots in Figure A.9. The posterior predictives encompass a majority of the experimental data within the 5 to 95% intervals. There are small discrepancies between the simulated realisations and experimental data that could indicate model misspecification [33]. For example, the maximum radius and filamentous area are on the boundary of the posterior predictive distributions at some time points. However, overall, the model is able to capture the experimental morphology for most summary statistics, with the posterior prediction residing closely around the true values of the summary statistics.

Figure 3.5 compares the marginal posterior distributions for the three strains. Pairplots of the posteriors can be seen in Appendix A.6. As anticipated, *n*^∗^ and *p_sp_* were the parameters with the most informative posteriors. For example, comparing AWRI 796 50 µm with AWRI 796 500 µm, we saw a big difference in *n*^∗^ which is expected as filamentation occurs earlier in the AWRI 796 50 µm than AWRI 796 500 µm due to the smaller initial nutrient level. Similarly, we expect *p_sp_* to be small as the experimental colonies have a limited number of distinct filaments. Since the number of filaments is controlled by the number of pseudohyphal cells emerging from sated cells, smaller *p_sp_* values will result in fewer filaments. Differences in *p_sp_* between different experiments can be explained by the amount of filamentation in each experiment. AWRI 796 50 µm has more filaments that AWRI 796 500 µm, and thus the AWRI 796 50 µm colony has a larger *p_sp_*. Therefore, although analysis of the synthetic data suggested that the NLE method may not recover the true value for *p_sp_,* it does identify relative changes in this parameter due to experimental conditions. These differences in *p_sp_* across experimental conditions were not detected using the ABC-based method and inference using data from a single time point [1].

**Figure 3.5:**
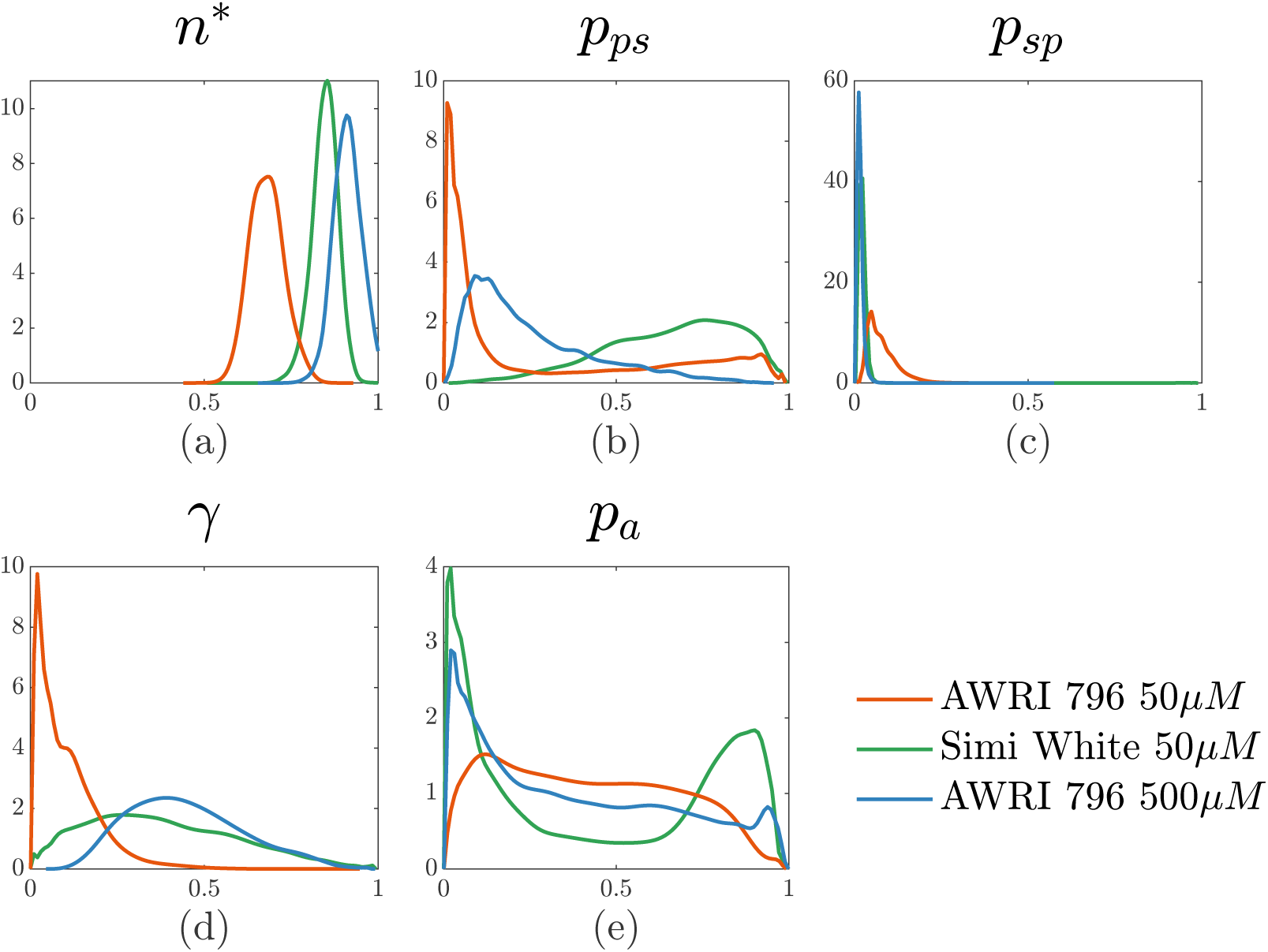
Comparison of the marginal posteriors for three experimental colonies. The colonies used here are AWRI 796 50 µm sample 1, AWRI 796 500 µm sample 8, and Simi White 50 µm sample 7.

**Figure 3.6:**
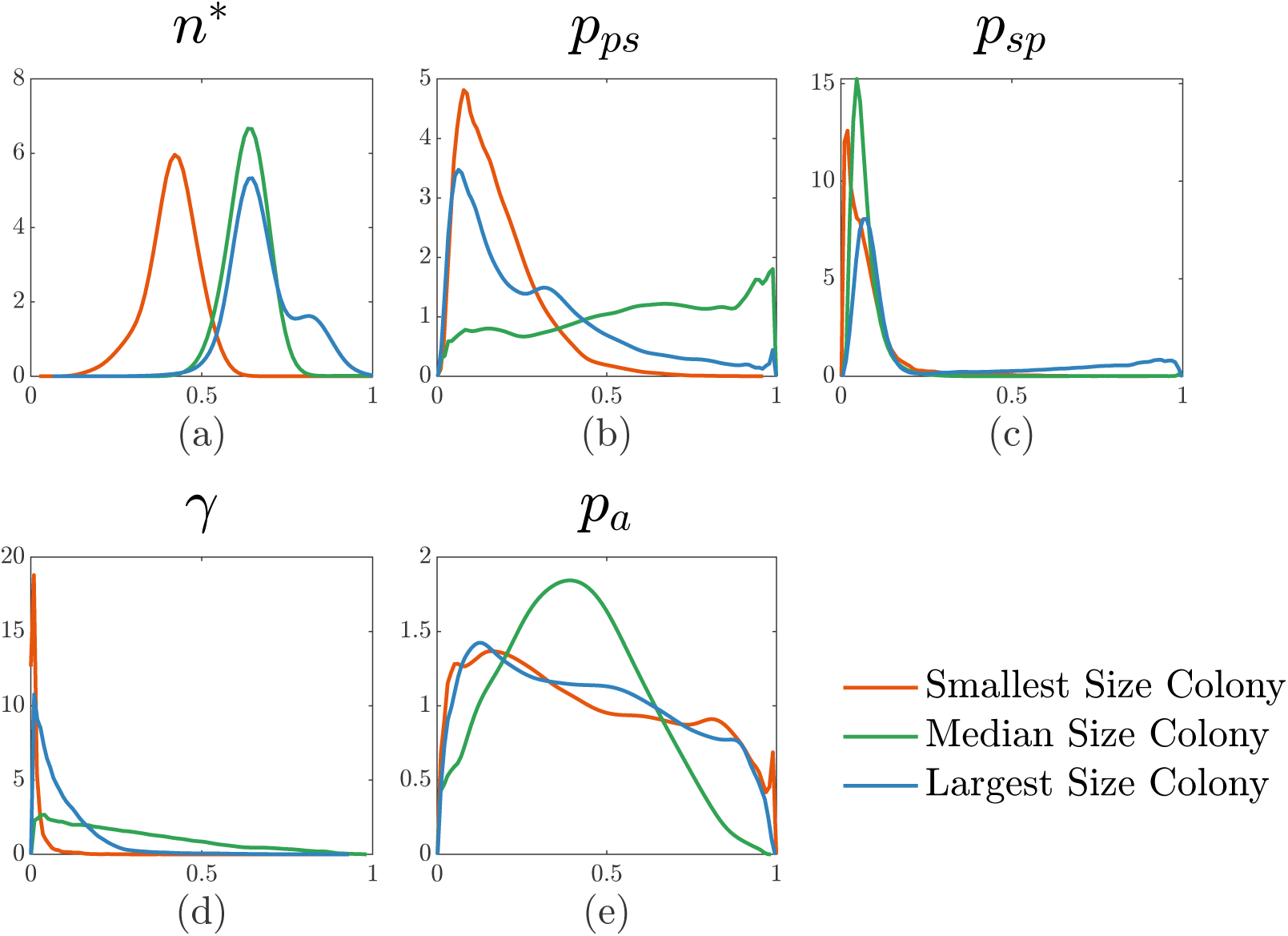
Comparison of posterior distributions of AWRI 796 50 µm colonies of different final areas. The smallest colony was sample 12, the median size was sample 14, and the largest was sample 9.

**Figure 3.7:**
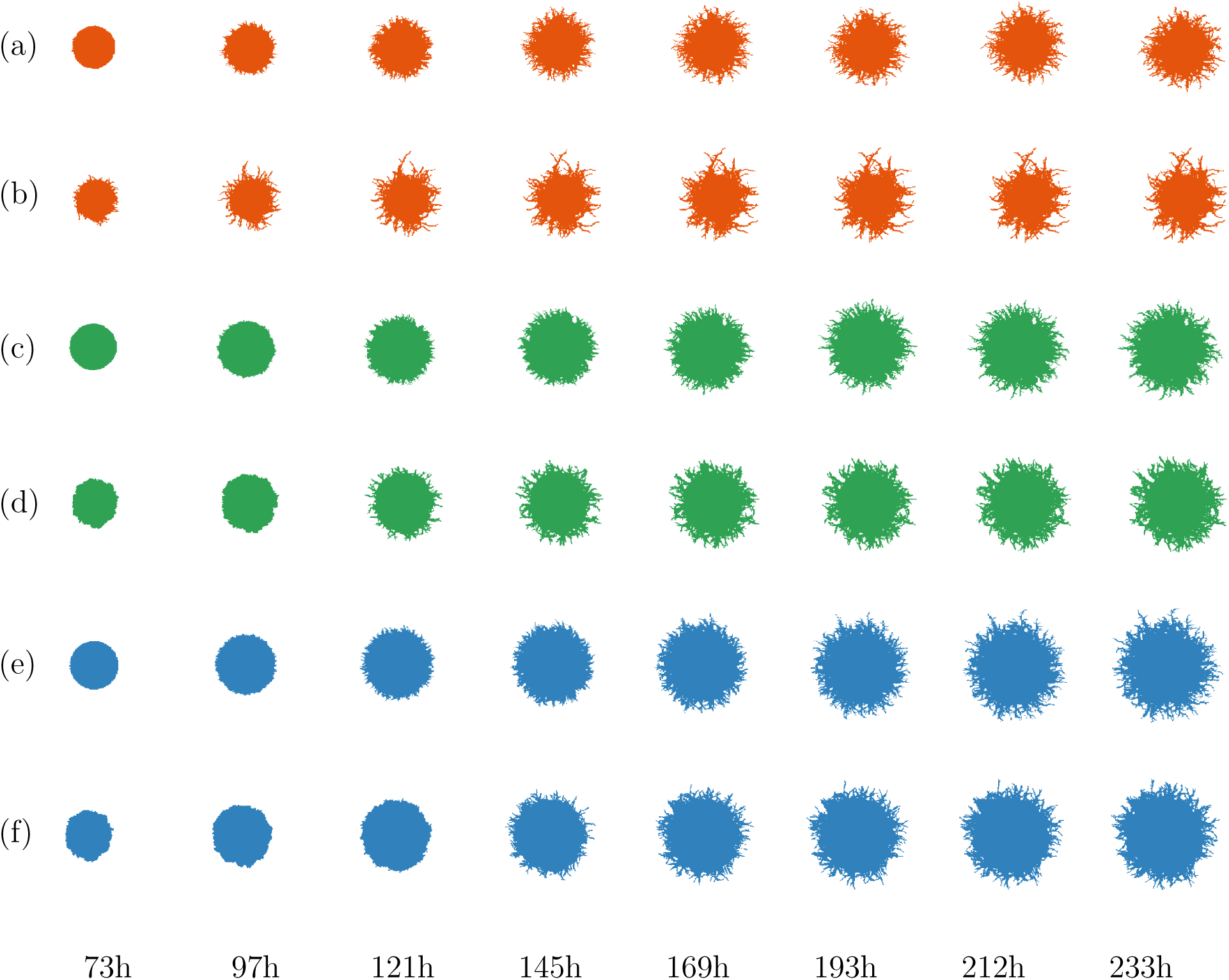
Simulation drawn from the posterior distribution in Figure 3.6 for different area colonies of AWRI 796 50 µm. (a) Sample 12, smallest size colony. (b) Simulation of Sample 12. (d) Experiment of Sample 14, medium-sized colony. (d) Simulation of sample 9 (e) Experiment of Sample 9, largest size colony. (f) Simulation of Sample 9.

**Figure 3.8:**
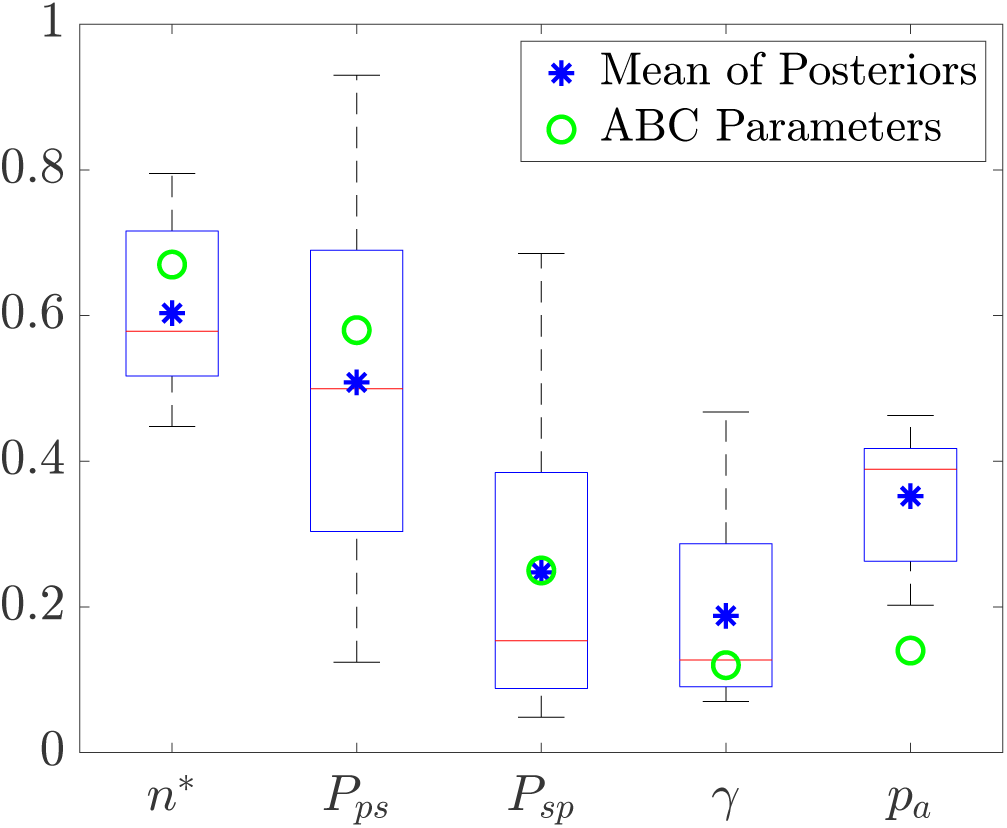
Boxplots derived from the means of the posterior distributions inferred from 14 experiments of AWRI 796 50 µm. The star represents the average of the means, the red bar represents the median value, and the circle markers represent the ABC mean parameters.

The posterior for the forking control parameter, *γ*, is wide for Simi White and AWRI 796 500 µm, but right-skewed for AWRI 796 50 µm. This can be explained by the small values of *p_sp_* for the Simi White and AWRI 796 500 µm, which limits the number of pseudohyphal cells being produced. Since forking only applies to pseudohyphal cells, with fewer pseudohyphal cells the effect of *γ* is diminished. When *p_sp_* takes larger values, such as in the AWRI 796 50 µm colonies with more filaments than other experiments, the forking parameter is more identifiable. Compared to Simi White 50 µm and AWRI 796 500 µm, the AWRI 796 50 µm colonies have more thin filament branches and increased forking. Therefore, the parameter *γ,* which limits forking, is lower for AWRI 796 50 µm than for the other two experimental conditions.

The posterior distribution for *p_ps_*, the probability of a pseudohyphal cell producing a sated cell, is difficult to interpret, suggesting that this parameter is unidentifiable. To test how *p_ps_* impacts morphology, we simulated colonies with varying the values of *p_ps_* in Appendix Figure A.13. These show that, at least for this combination of parameters, changing *p_ps_* had minimal impact on the overall colony morphology. Another uninformative parameter is *p_a_*. This parameter controls the probability of choosing a sated cell to proliferate over a pseudohyphal cell after filamentation has been triggered. Hence, these two parameters *p_ps_* and *p_a_* affect the thickness of the filaments of the simulated colonies. However, currently our summary statistics do not capture the filament thickness, which could be the reason why these parameters are uninformative.

### 3.3 Variability within replicates of the same experimental conditions

Due to large variability in final colony area across replicates with the same experimental conditions (see Appendix Figure A.18), we also investigated the impact of colony area on the inferred posteriors. We selected three colonies with the smallest, median and maximum area from the AWRI 796 50 µm experiments and performed inference on these. We observed from the posterior distributions (see Figure 3.6) that different areas do impact our estimates of the parameters for our model. The mean value of *n*^∗^ increases as the size of the colony increases. Moreover, the distributions of *p_sp_* and *γ* are similar between colonies while *p_sp_* and *p_a_* are uninformative.

In Figure A.17, we present the prior and posterior predictive distributions. For the smallest colony, the mean radius, branch length and compactness lie within the 25–75% prediction intervals, while the maximum radius and filamentous area are less accurately predicted. As previously noted, this could be due to model misspecification. However, an interesting trend is that the uncertainty in the predictive intervals decreases for the larger colonies, with the intervals being more centred around the experimental summary statistics (black lines). We also visualise colonies drawn from the posterior distributions as shown in Figure 3.7. As reflected in the predictive distributions, the smallest colonies appear the least similar to their experimental counterparts, while the simulations match best for the largest colony.

To further investigate the variability in parameter values within a set of experimental conditions, we performed inference on all 14 individual colonies of AWRI 796 50 µm. We then took the mean of each approximate posterior and compared these point estimates with those derived using ABC by Li et al. [1]. Box plots of the means are shown in Figure 3.8 along with the ABC results. The most impactful parameter on colony morphology, *n*^∗^, is similar to the value observed in the ABC inference. The variability in this parameter is also small, as expected. Similarly, the other parameters are in good agreement. It is important to note that the variability in the parameter values is expected and is due to the large variability of colony area across replicates of the same experimental conditions, as shown in Appendix A.8. In addition, we do not expect the two estimates of the parameter means to be the same due to different techniques being used, and the different priors used. That is, in the ABC, due to computational complexity, we chose weakly informative priors, whereas in the NLE, we used uniform priors. However, as both techniques gave similar results, it suggests the assumptions made by Li et al.

In this work, we have been comparing independent inferences that assume the parameters are different within the same experiment. We also investigated using a joint posterior approach to combine individual colony parameters within an experimental condition. This approach assumes that the parameter values are the same within a set of replicates of the same experimental condition. From the posterior predictive distribution, we found that the joint posterior approach performed poorly at capturing the colony morphologies compared to the individual inference shown in Figure A.17. This suggests that colonies within the same set of experiments are subject to random effects that change the individual parameters for each colony.

## 4 Discussion and Conclusion

We used neural likelihood estimation and a time series of photographs to infer parameters of an agent-based model for filamentous yeast colonies. We first verified our inference on synthetic data and found that using a full time series of 48 summary statistics (8 photographs, 6 summary statistics per photograph) provided the most informative posterior distributions. We showed our model could successfully capture the morphology of colonies grown in three experimental conditions. The most important determinants of colony morphology were *n*^∗^, the threshold size for filamentous growth, and *p_sp_,* the probability of a sated cell producing a pseudohyphal cell.

We used a basic neural likelihood estimation (NLE) rather than sequential neural likelihood estimation (SNLE) that trains the model over a number of rounds. Initial testing using SNLE was performed where convergence was determined using the Euclidean distance between simulated (synthetic) parameter values and means of the posterior distributions. Convergence happened within a single round. Most likely, this was due to a relatively large (1,000 data points) sample size. However, if our sample size was small (100 data points), then our results would have benefited from SNLE. Another added benefit of using NLE was that simulation and inference took roughly one day compared to weeks for the ABC method used by Li et al. [1]. NLE also allows us to reuse simulations for independent colonies and run them up to specific areas, compute respective summary statistics, and thereby further reduce computational costs. Furthermore, the analysis in Section 3.3 required training 14 different models. Although this was not as computationally expensive as it first seems because some data could be reused, it would be better to train a single model that also takes the area as a parameter. Additionally, a different likelihood model, such as an autoregressive conditional density model [34], may better capture the dependencies in the time-series data than the Gaussian mixture model.

Some parameters of our model were unidentifiable, suggesting the possibility of simplifying the model to have fewer parameters while maintaining the ability to produce similar morphologies. Alternatively, we would need to add new summary statistics to better capture the aspects of the morphology controlled by these parameters. For example, the current statistics lack the ability to capture branch thickness. One potential improvement could be to use a hierarchical random effects model [35], which is most appropriate for analysis of multiple colonies from the same experiment. Fitting such a model is beyond the scope of this work, as it requires properly accounting for time and differences in initial conditions.

There were small discrepancies between the predicted and experimental colonies, perhaps due to model misspecification. This possible misspecification suggests that we can make improvements to our technique, such as including a summary statistic that quantifies the thickness of individual filaments and an angular metric to quantify the spread of the filaments [31]. Future work could include developing a three-dimensional model of filamentous growth with growth in the *z*-direction invading the agar. Cells adhering to and invading the agar medium are of great interest to biologists because of their importance in pathogenic infections.

## Acknowledgements

KL acknowledges funding from the Australian Government through a Research Training Programme Scholarship. BJB, JEFG, AKYT, and VJ acknowledge funding from the Australian Research Council (Grant numbers DP230100406, DE240100097). This work was supported with supercomputing resources provided by the maths1 High Performance Computing service at the University of Adelaide.

## Data, Code and Materials

All the data, code and materials can be found on GitHub.

## A Supplementary Material

### A.1 Logit transform of parameters

When sampling out the posterior, we include a Jacobian term in the priors because of the logit transformation performed on the parameters *θ*. Beginning with the logit transform,

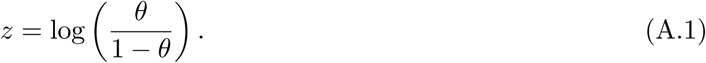

The inverse logit transform of Equation (A.1) is

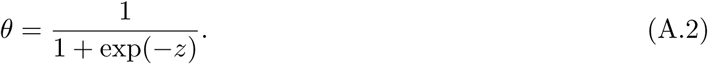

The probability density function, *P* (*θ*) of the Beta distribution defined on a unit interval is given as

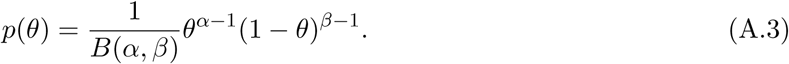

When we transform *θ* by applying the logit transform, we need to account for the change using the derivative d*θ/*d*z*. Using the transformation of the variable, the transformed density becomes,

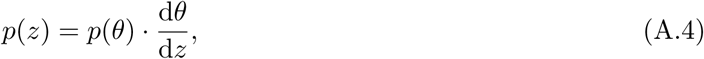

where d*θ/*d*z* is the Jacobian. Taking the derivative of Equation (A.2) with respect to *θ* we get,

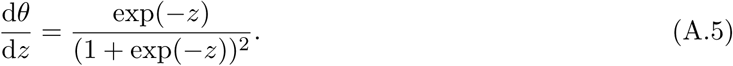

Substituting Equations (A.3) and (A.5) into Equation (A.4) to get the transformed density,

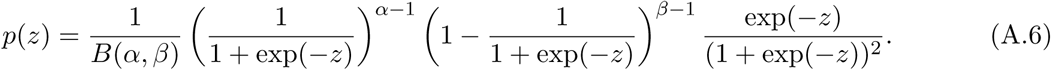

### A.2 Synthetic data

**Figure S1:**
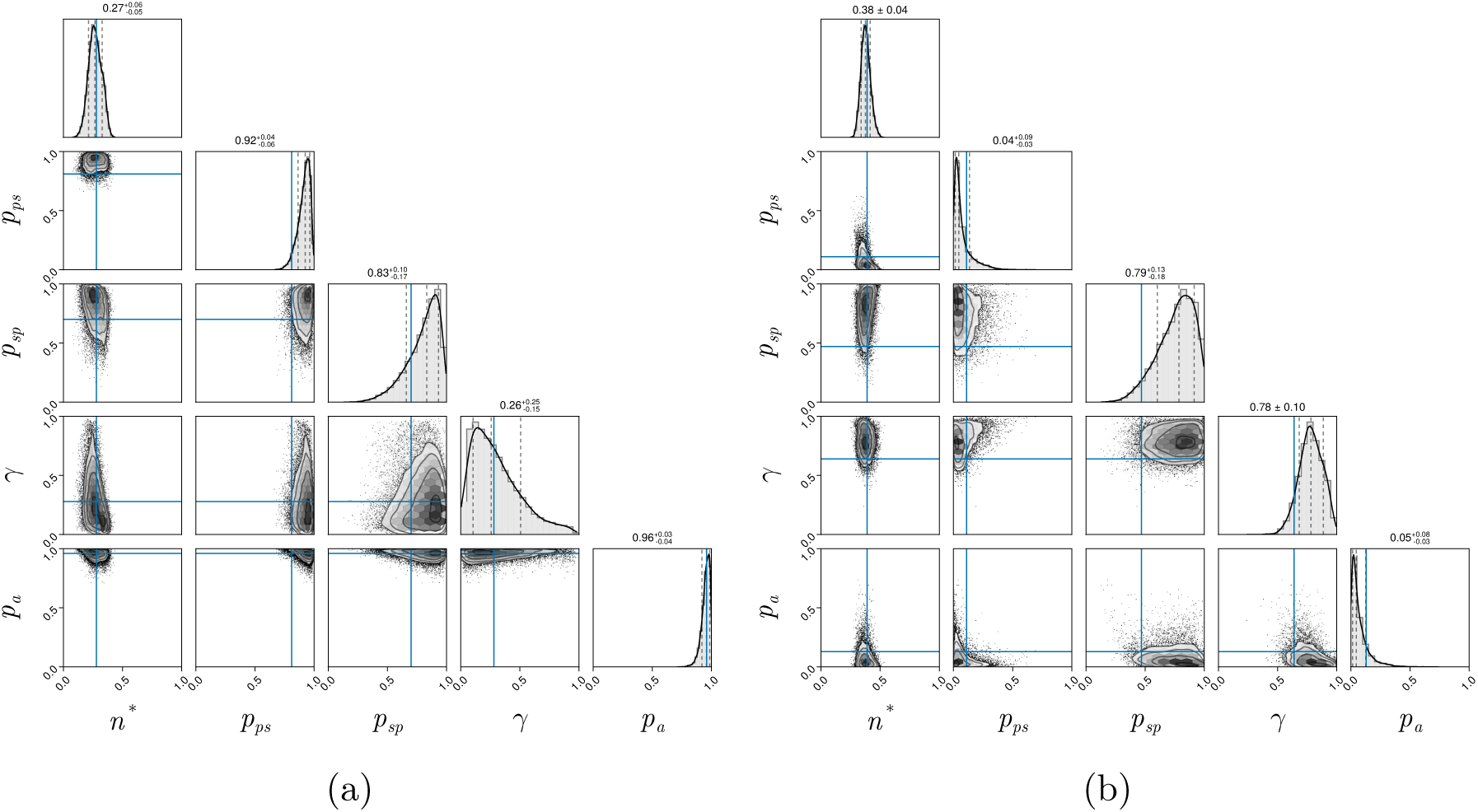
Pair plots of synthetic posterior distribution. The blue line represents the ground truth parameter values. (a) *n*^∗^ = 0.28, *p_ps_* = 0.81, *p_sp_* = 0.70, *γ* = 0.28, *p_a_* = 0.96. (b) *n*^∗^ = 0.39, *p_ps_* = 0.11, *p_sp_* = 0.47, *γ* = 0.64, *p_a_* = 0.13.

**Figure S2:**
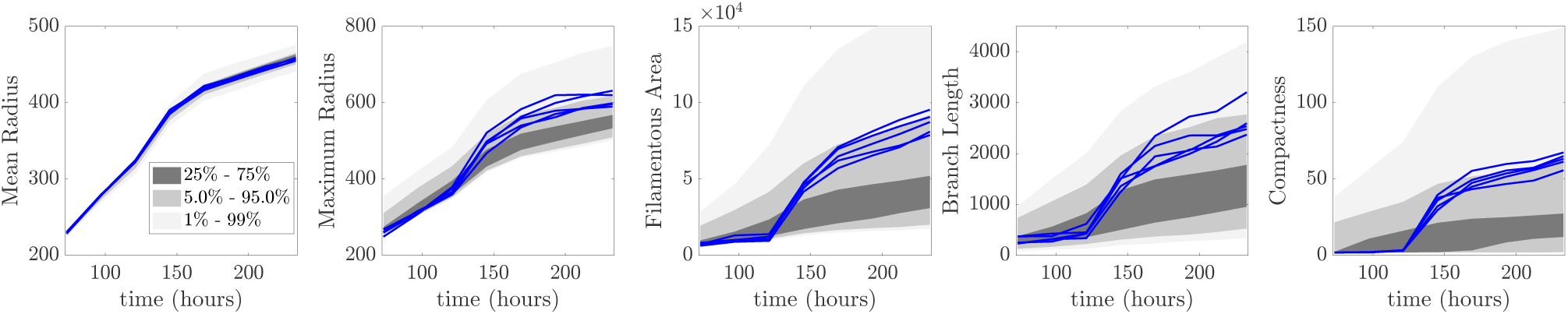
Summary statistics from simulations of the same parameters simulated up to the same area. This shows the variability within a simulation for a fixed set of parameters.

We sample parameters from the posterior of the synthetic dataset to show the variability in colony morphology of ten realisations. The parameter values are reported in the left corner of each simulation with the ground truth in panel (a). As *n*^∗^ is the most important parameter in our model, we vary the ground truth colony *n*^∗^ from *n*^∗^ = 0.28 (Figure A.4), *n*^∗^ = 0.39 (Figure A.5), to *n*^∗^ = 0.67 (Figure A.6). All three sets of simulated results sampled from the posterior align well with the ground truth.

**Figure S3:**
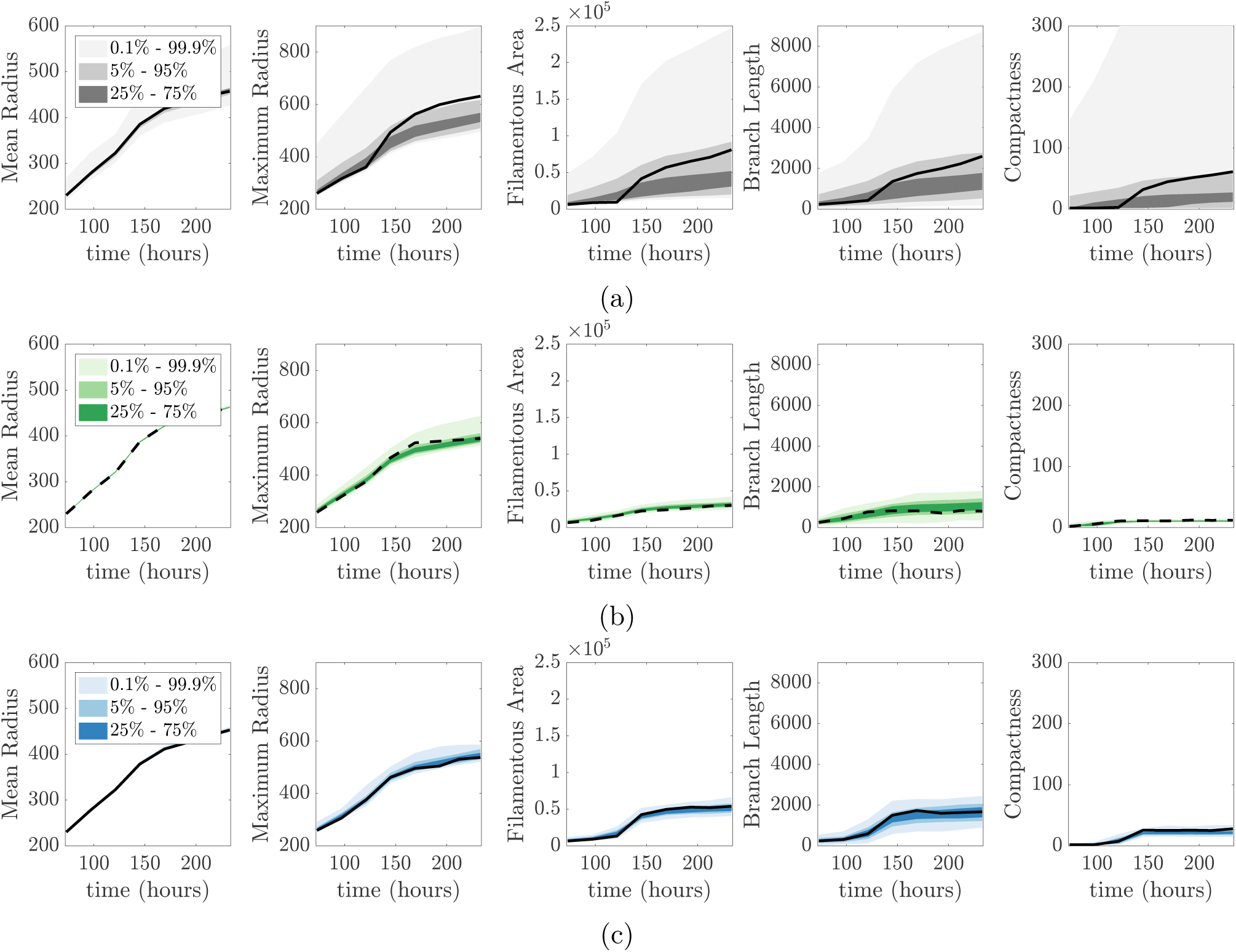
Predictive distribution for the synthetic datasets. (a) Prior predictive distribution with the black lines representing the experimental data. The following are the posterior predictive distributions with parameters (b) *n*^∗^ = 0.28, *p_ps_* = 0.81, *p_sp_* = 0.70, *γ* = 0.28, *p_a_* = 0.96. (c) *n*^∗^ = 0.39, *p_ps_* = 0.11, *p_sp_* = 0.47, *γ* = 0.64, *p_a_* = 0.13.

**Figure S4:**
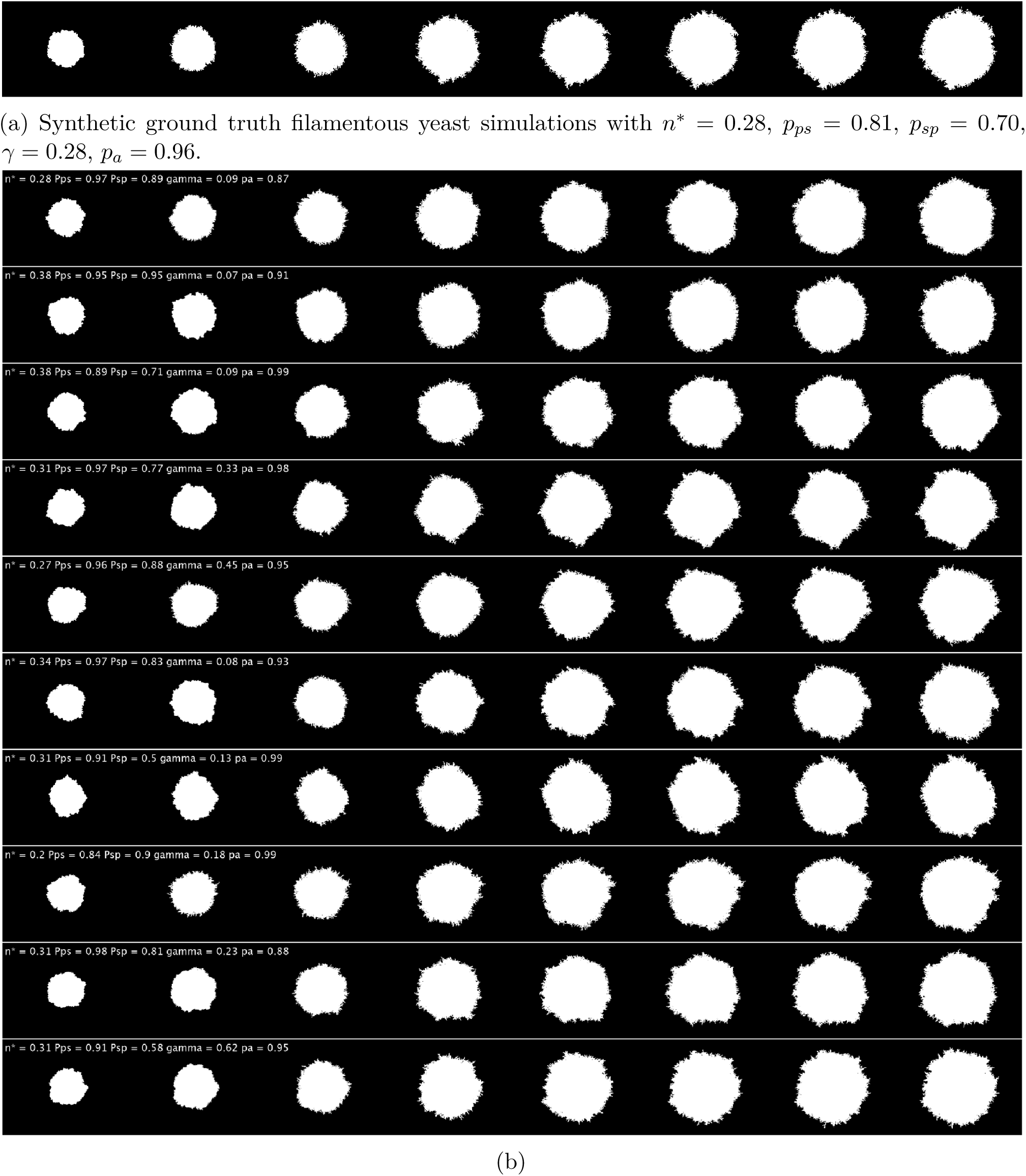
Colony simulation comparison between (a) Ground truth and (b) NLE prediction of ten realisations.

**Figure S5:**
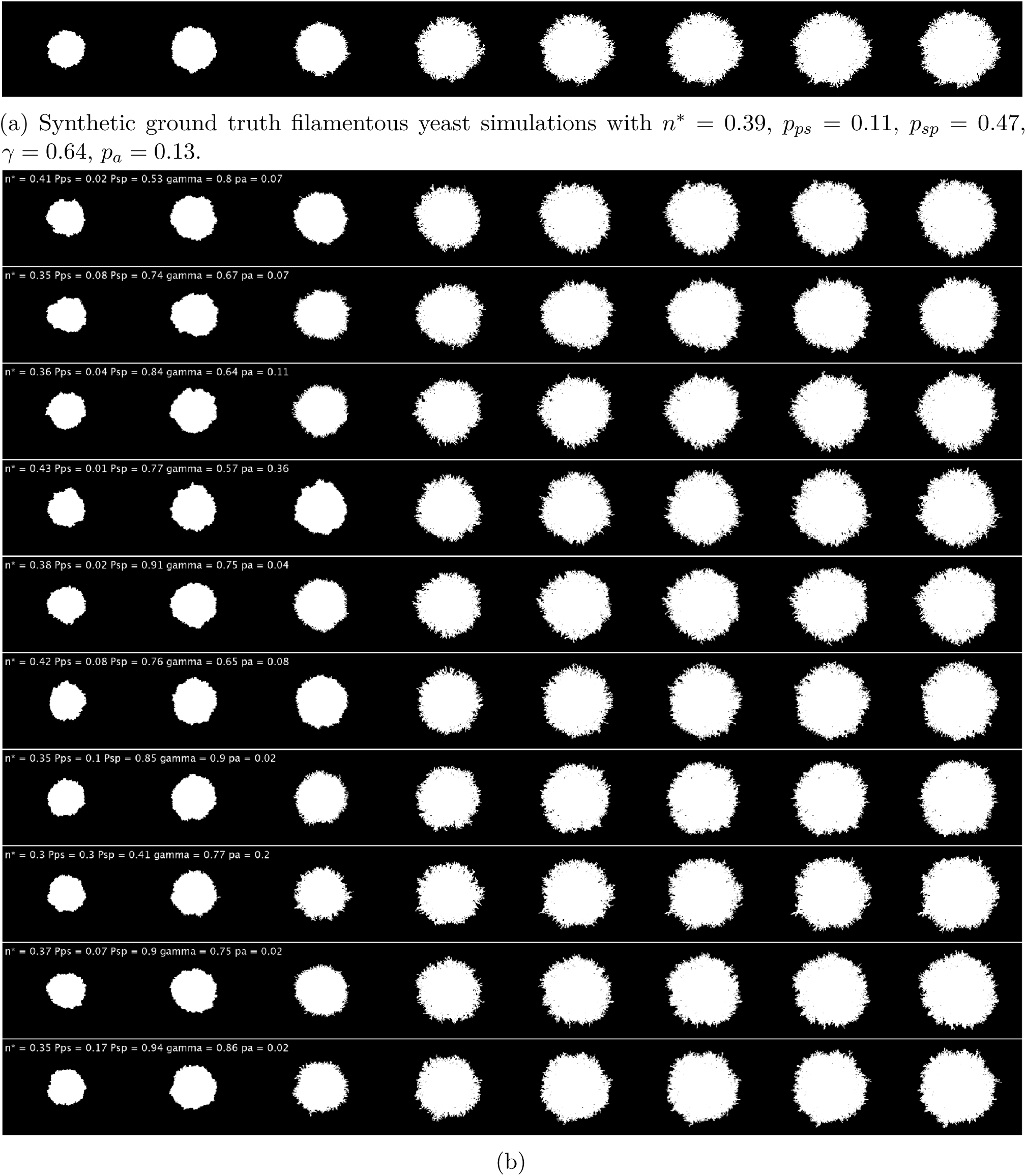
Colony simulation comparison between (a) Ground truth and (b) NLE prediction of ten realisations.

**Figure S6:**
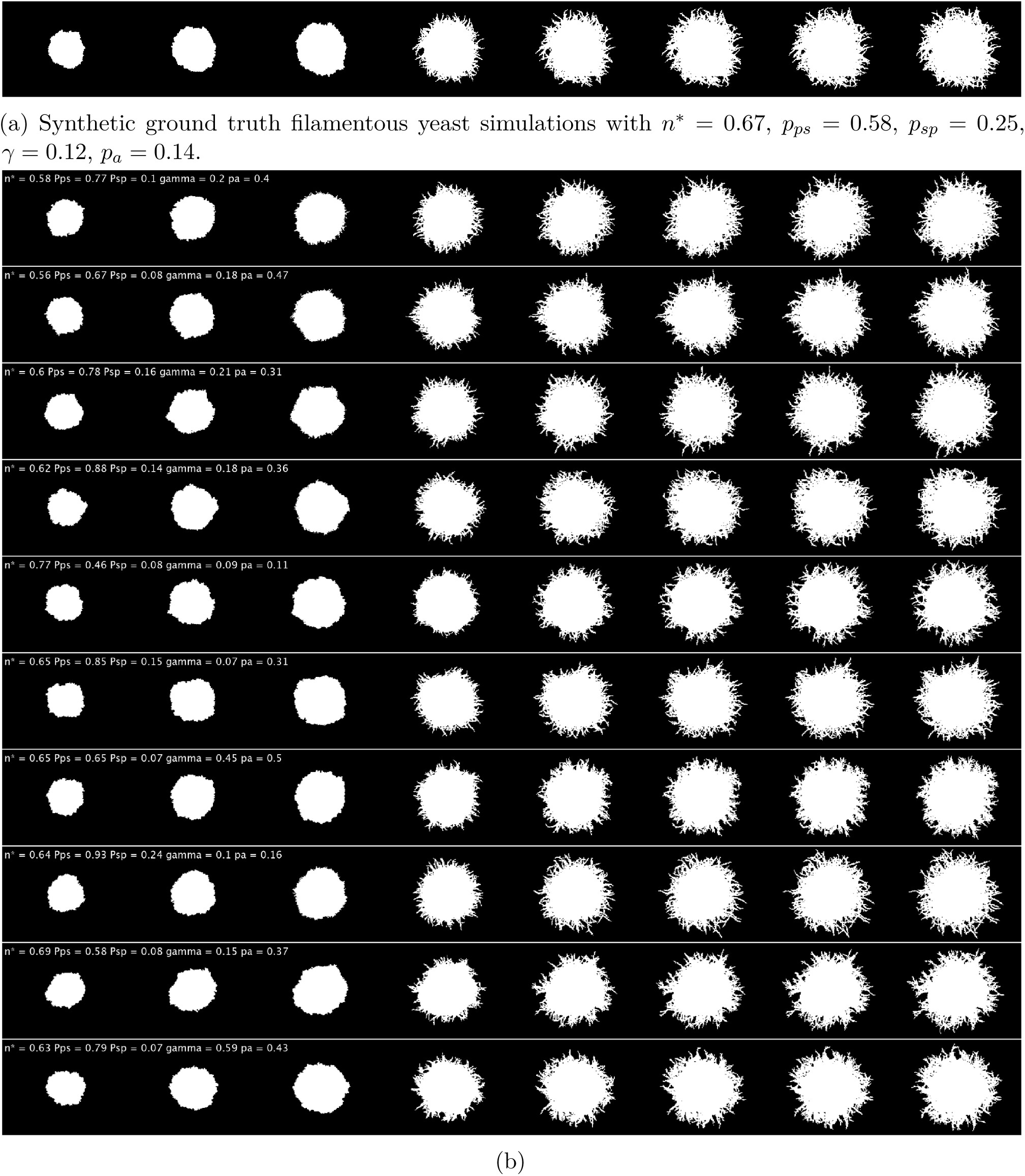
Colony simulation comparison between (a) Ground truth and (b) NLE prediction of ten realisations.

### A.3 Synthetic data using different numbers of summary statistics

**Figure S7:**
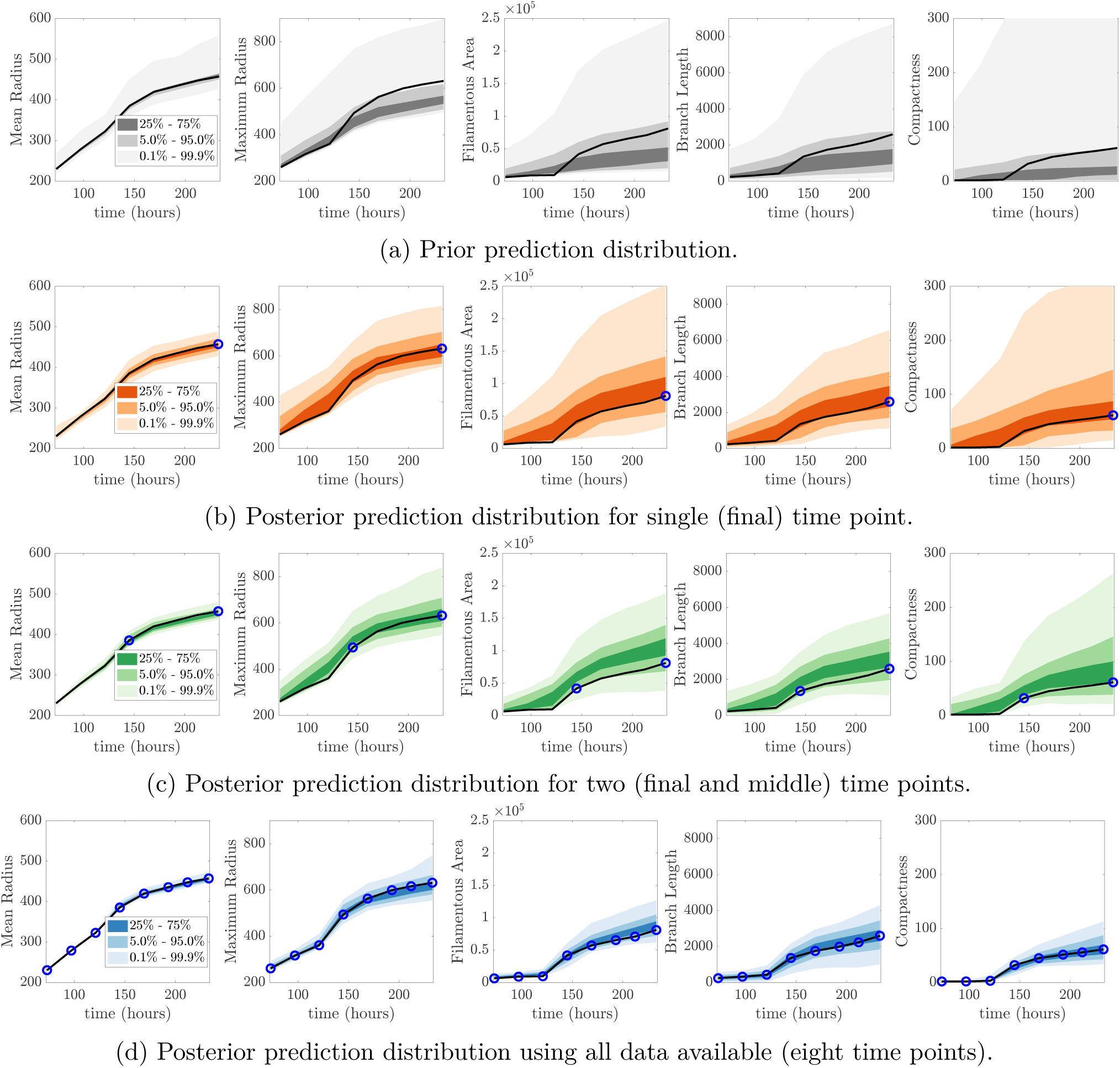
(a) Prior predictive distribution. Posterior predictive check for (b) eight time points, (c) two time points and (d) a single time point. The blue markers indicate the time points where we evaluate the summary statistics. (b,c,d) share the same prior predictive distribution shown in (a).

**Figure S8:**
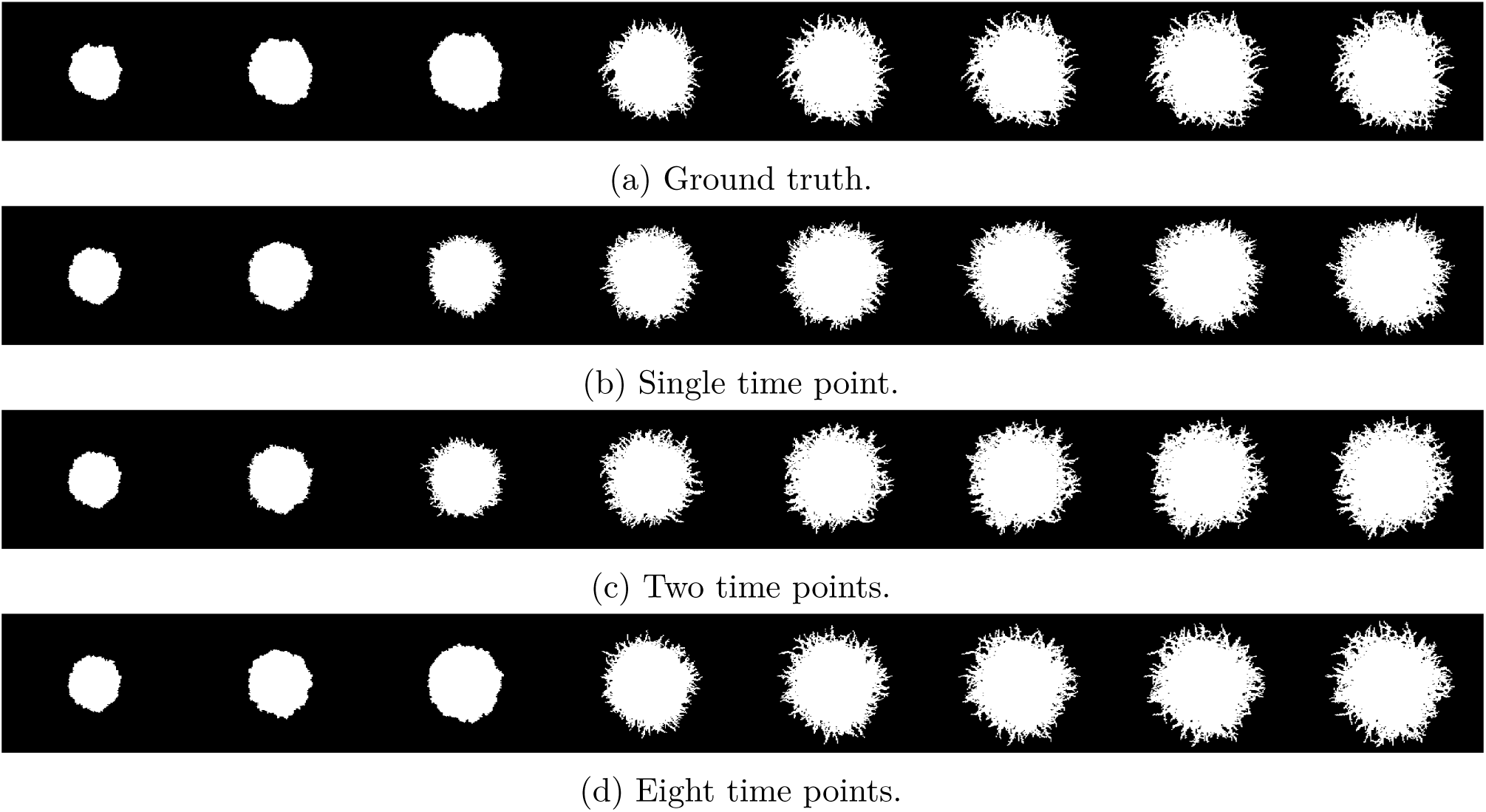
Comparison of simulations generated from posterior distributions inferred using different numbers of summary statistics. (a) The ground truth is plotted for comparison against a colony inferred from (b) a single time point (final time), (c) two time points (middle and final time) and (d) all time data available (eight time points).

### A.4 Experimental data

Here we present the experimental data, predictive distribution, and ten realisations drawn from the posterior distribution.

**Figure S9:**
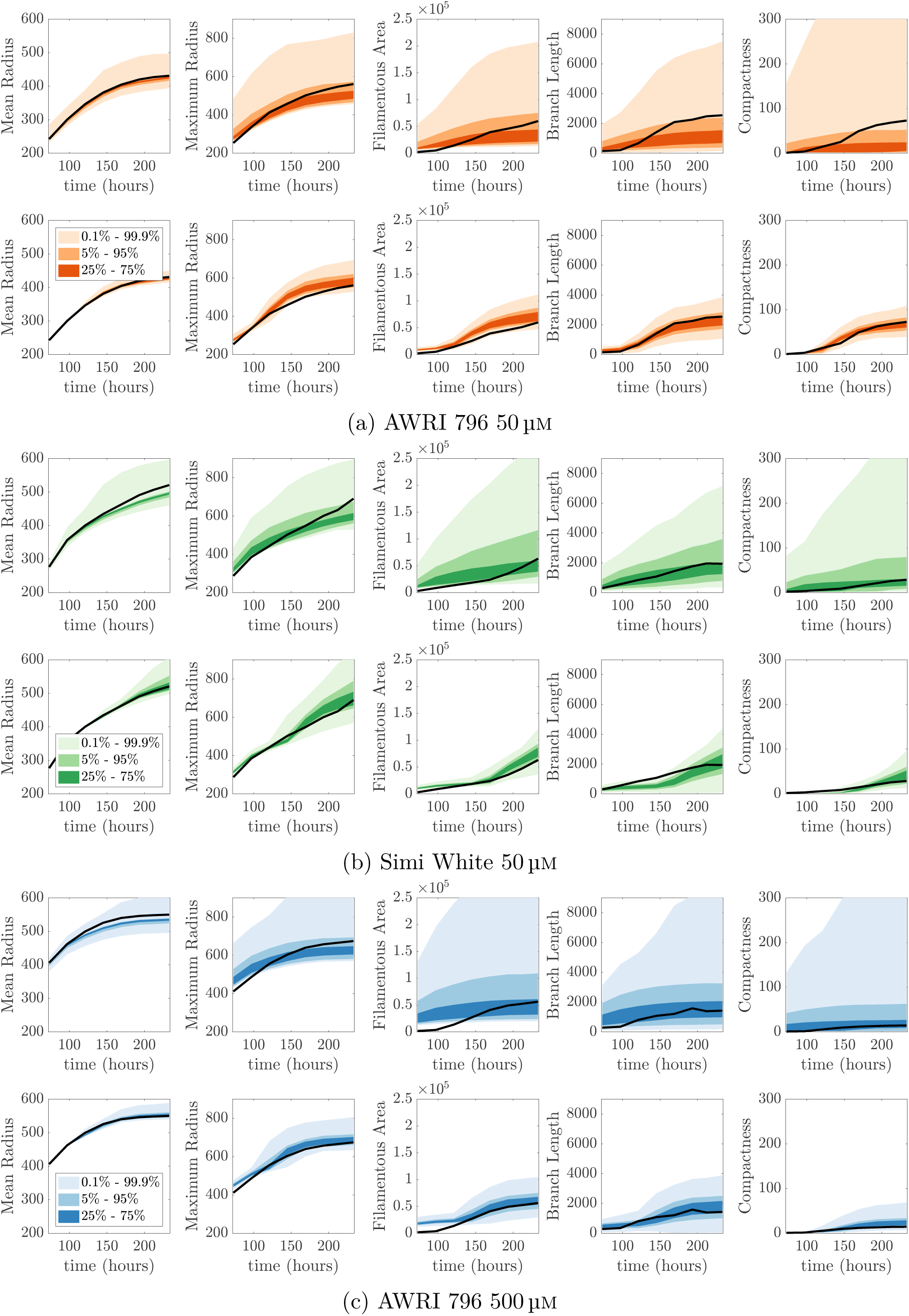
Prior predictive plots (top row) and their corresponding posterior predictive distribution (bottom row) for each experimental colony. (a) AWRI 796 50 µm (b) Simi White 50 µm and (c) AWRI 796 500 µm.

**Figure S10:**
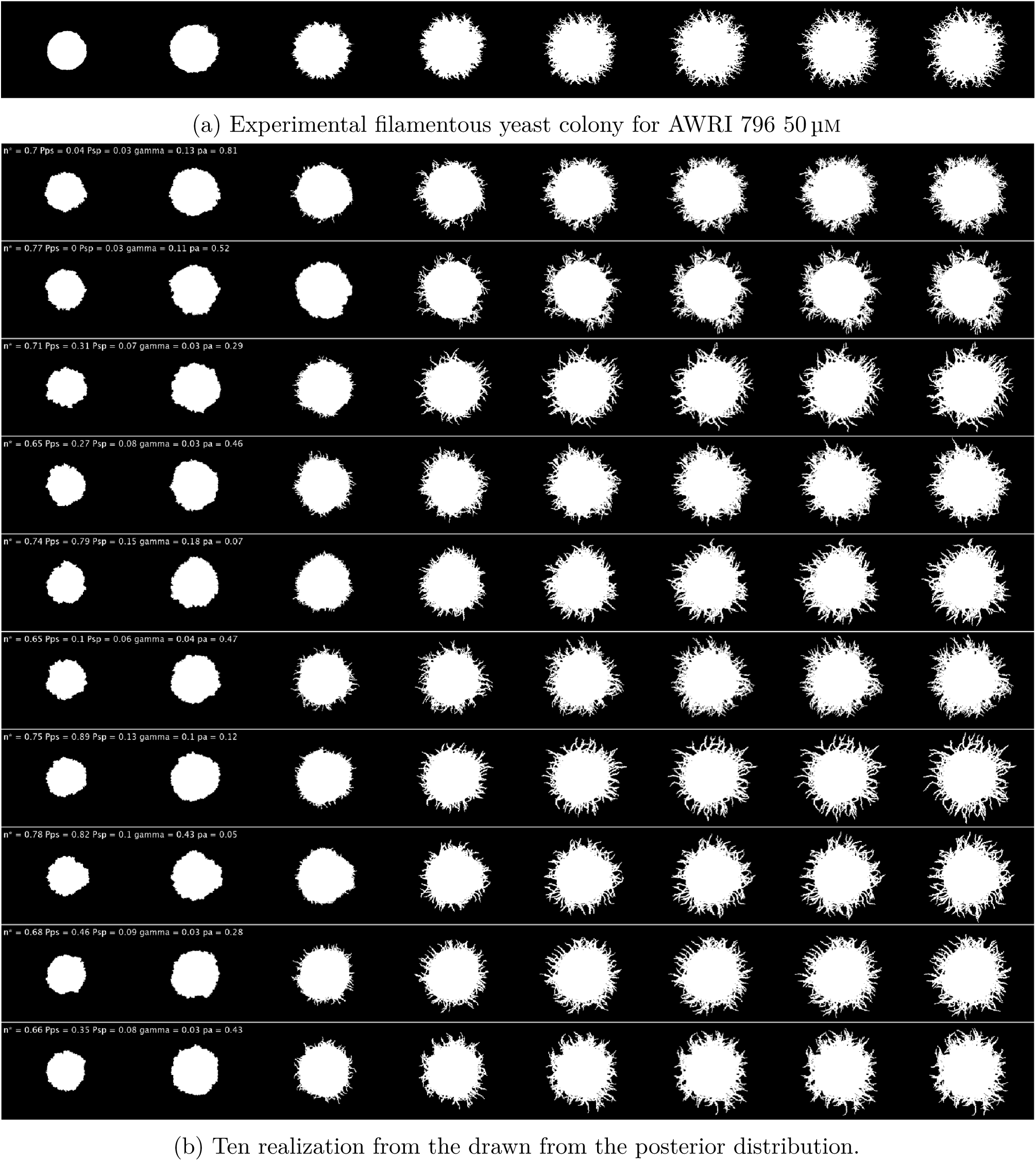
Colony comparison between (a) experimental AWRI 796 50 µm and (b) ten realisations drawn from the posterior distribution.

**Figure S11:**
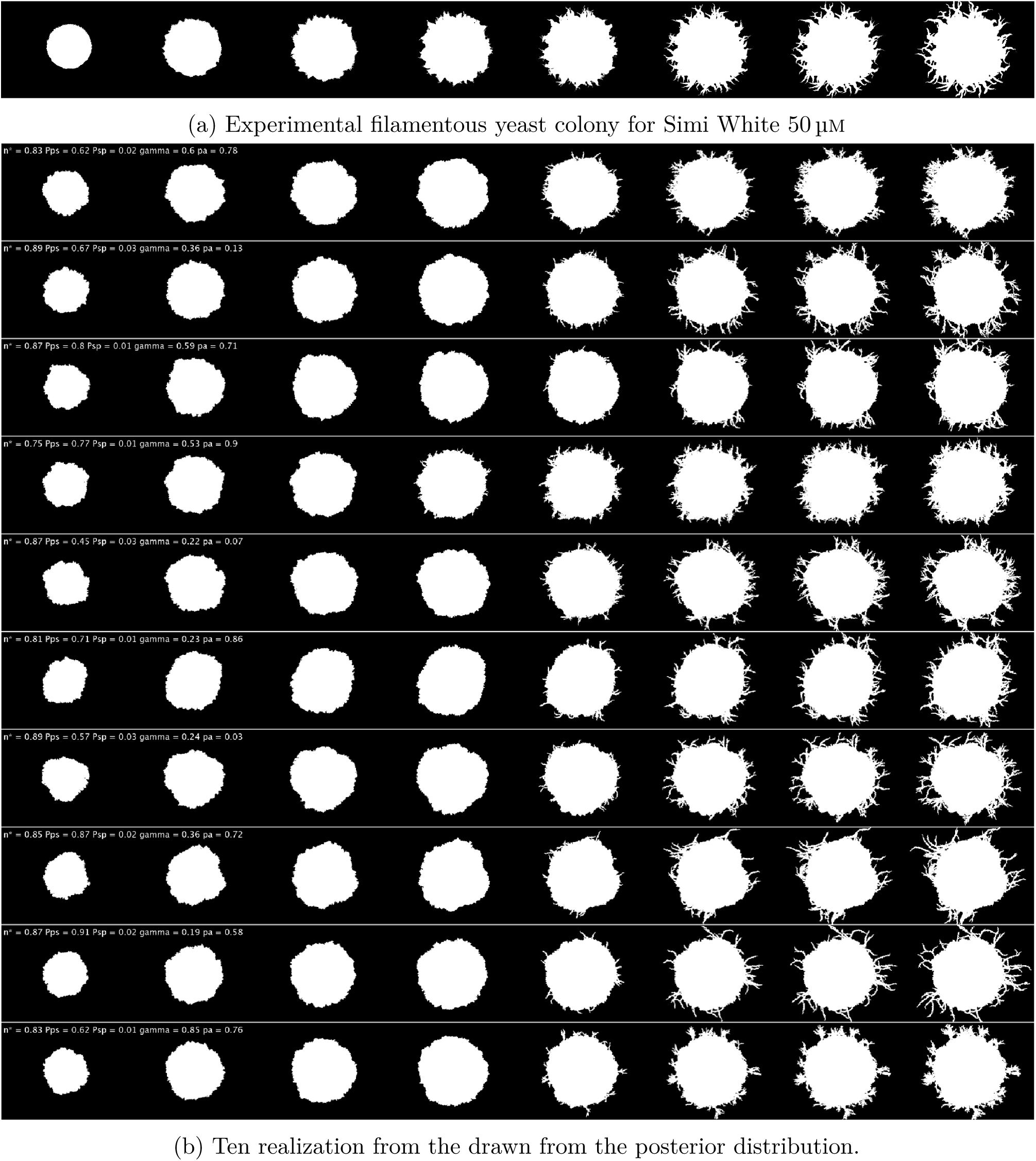
Colony comparison between (a) experimental Simi White 50 µm and (b) ten realisations drawn from the posterior distribution.

**Figure S12:**
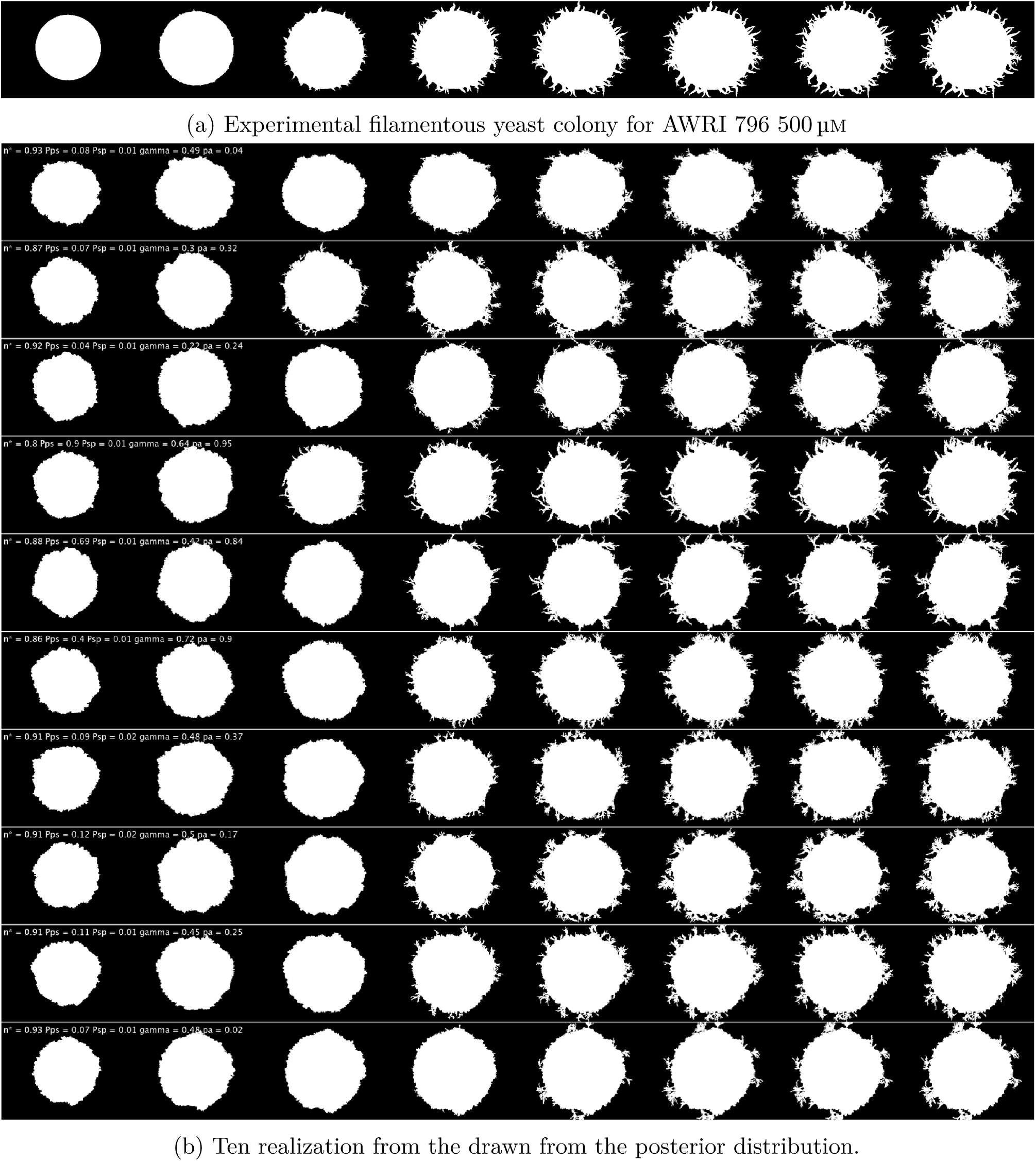
Colony comparison between (a) experimental AWRI 796 500 µm and (b) ten realisations drawn from the posterior distribution.

### A.5 Testing the impact of ***p_ps_*** on colony morphology

To demonstrate that the parameter *p_ps_* is uninformative, we fixed all other parameters constant and varied *p_ps_* from 0.1 by increments of 0.1 up to 1 and simulated the associated colonies presented in Figure A.13. As expected, we see the simulated colony morphology align closely with the experimental colony, with very little difference between simulations of varying values of *p_ps_*.

**Figure S13:**
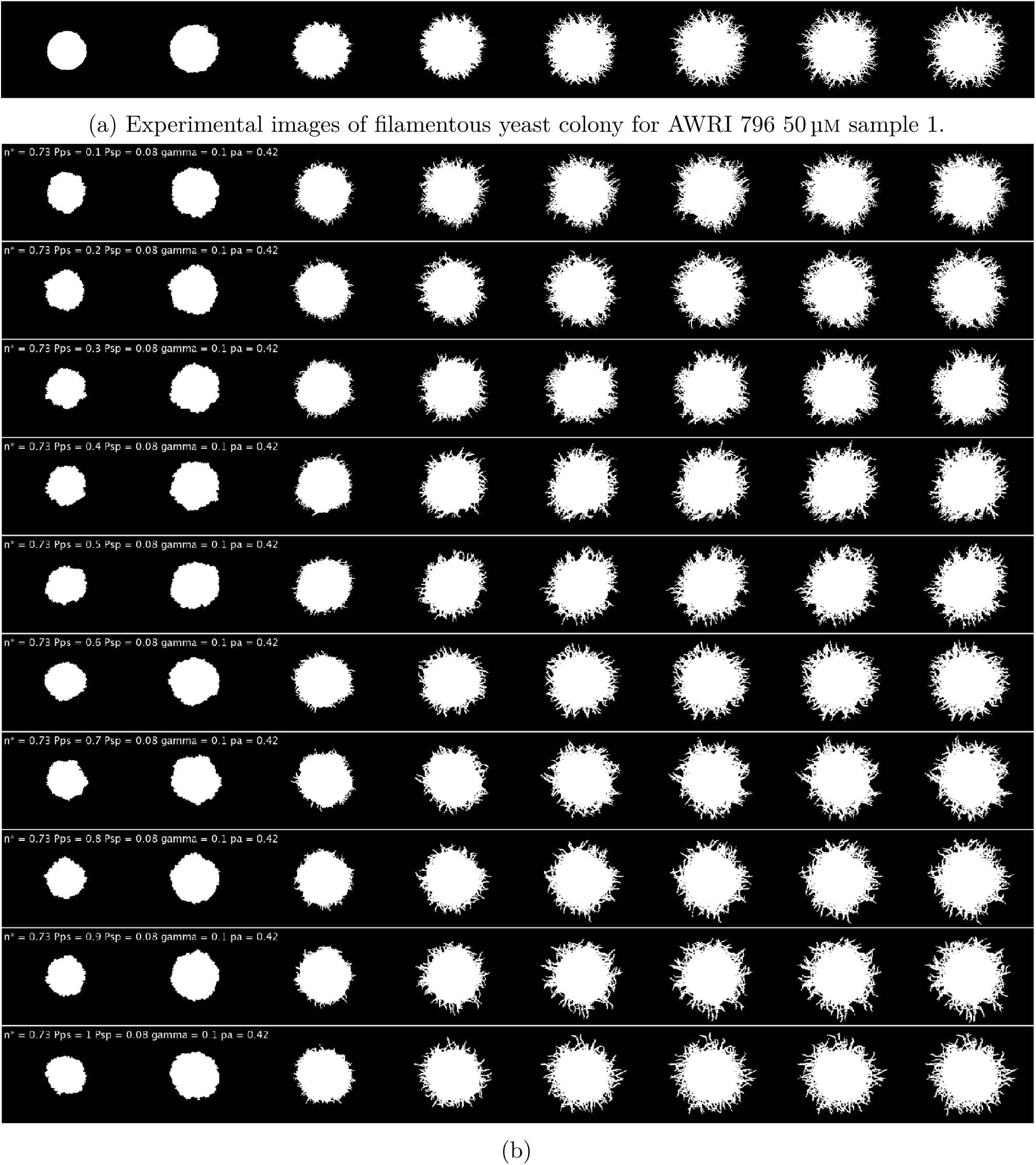
Colony comparison between (a) experiment of AWRI 796 50 µm and (b) each realisation is a different value of *p_ps_* while hold all other parameters constant (values reported in Figure 3.5) demonstrating this parameter is non-informative.

### A.6 Pairplots of experimental posterior distributions

Pairplot of posteriors of experimental colonies for AWRI 796 50 µm (Figure A.14), Simi White 50 µm (Figure A.15) and AWRI 796 500 µm (Figure A.16).

**Figure S14:**
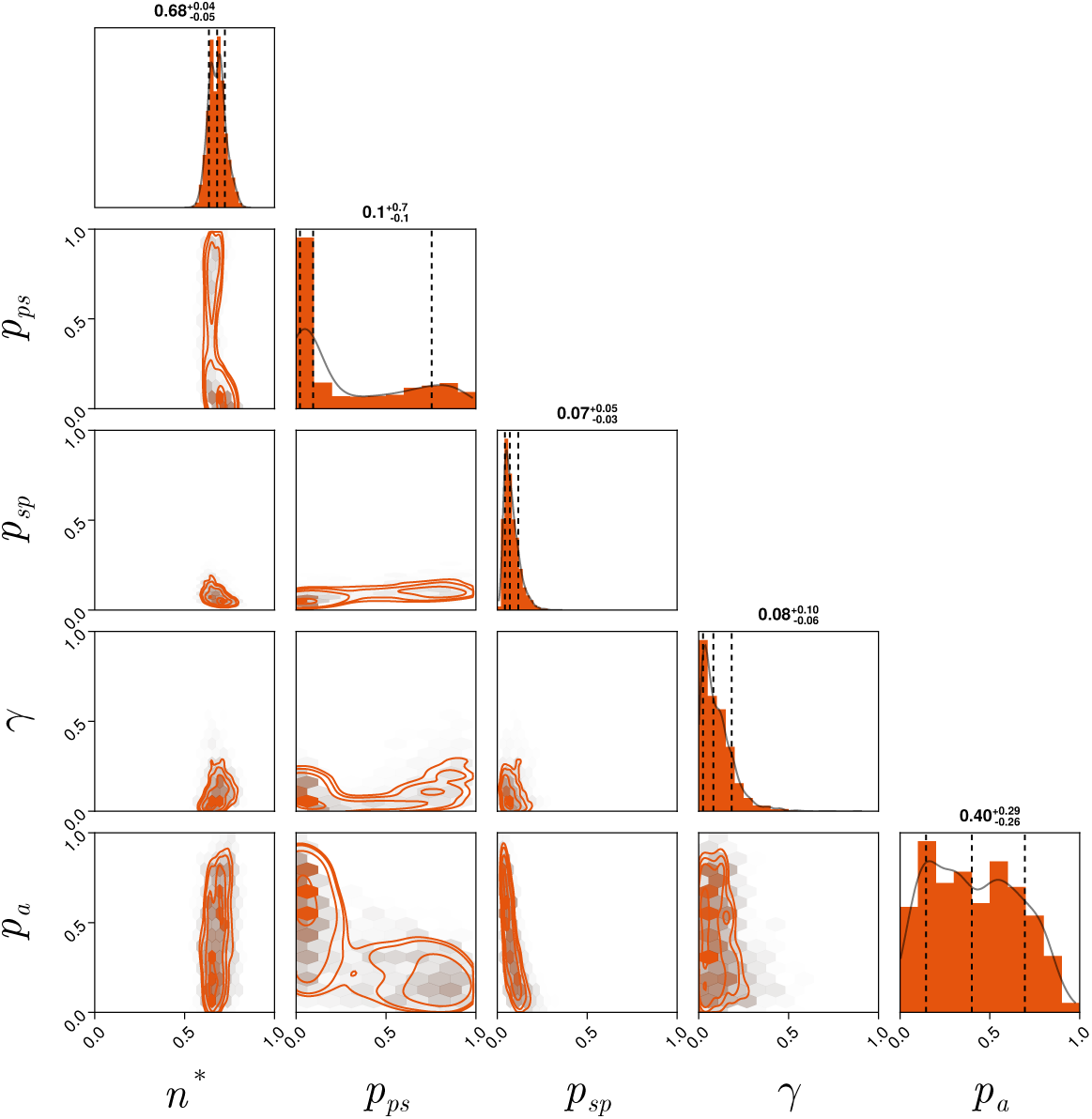
Pair plot of AWRI 796 50 µm posterior distribution.

**Figure S15:**
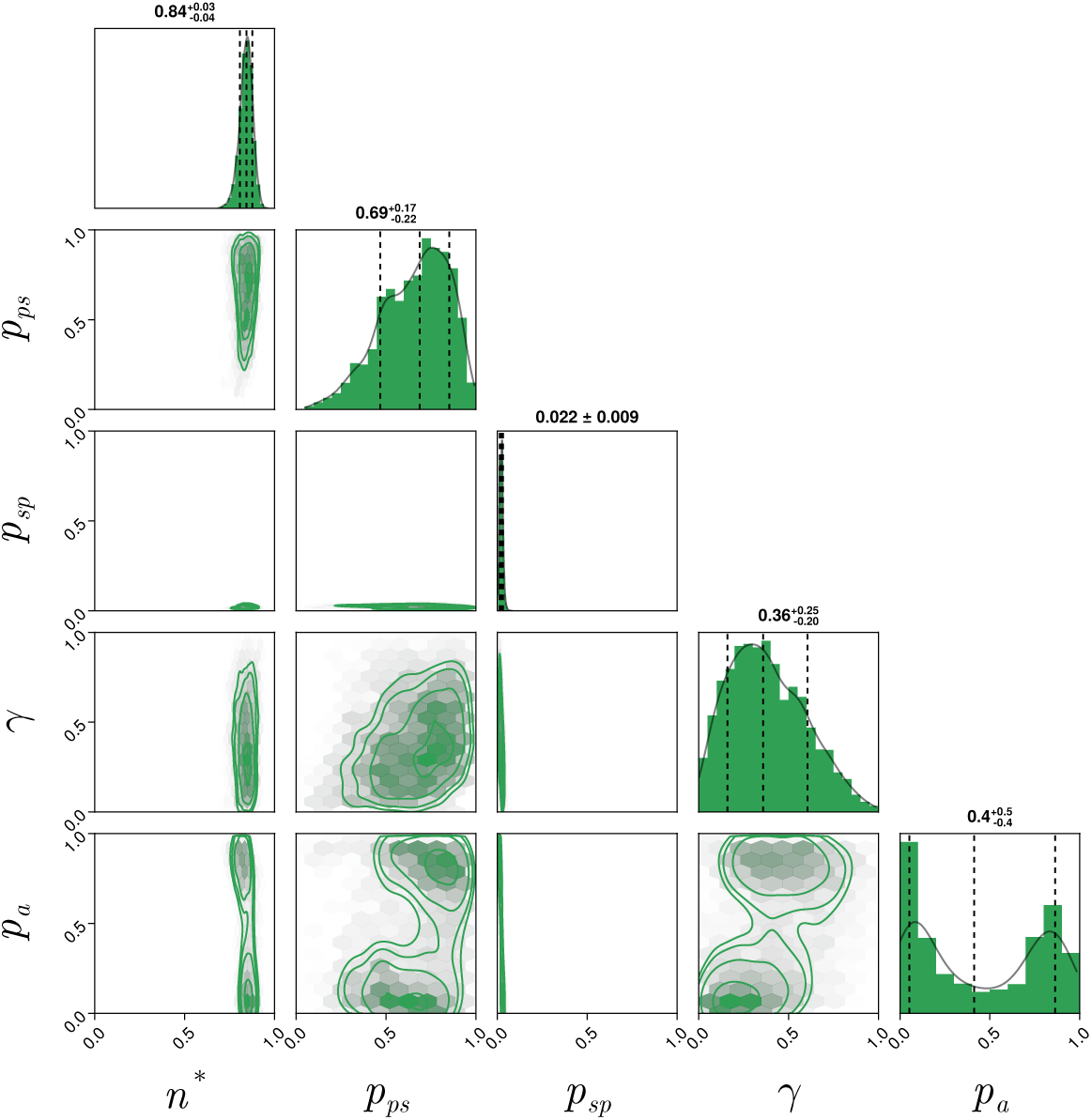
Pair plot of Simi White 50 µm approximate posterior distribution.

**Figure S16:**
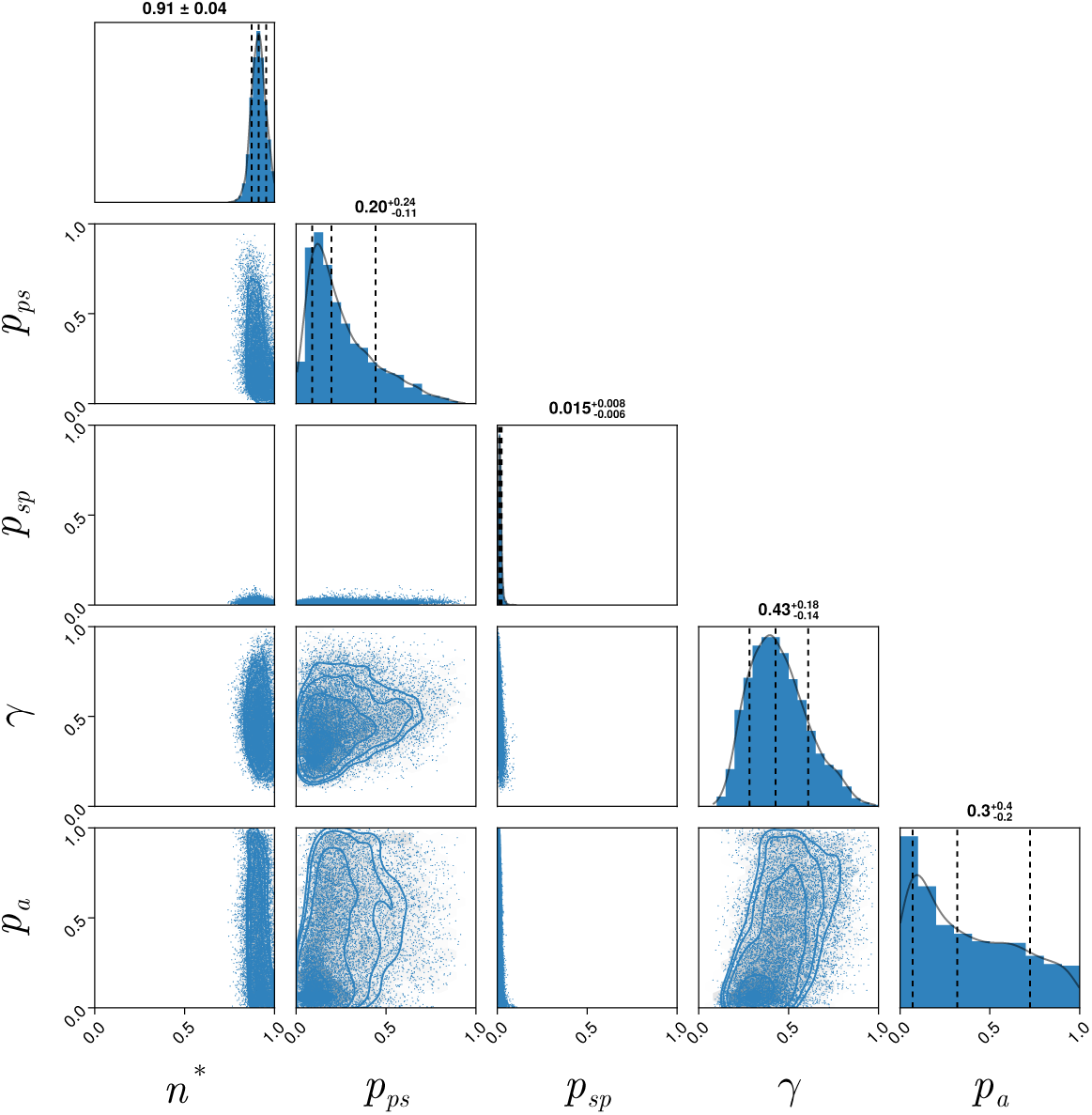
Pair plot of AWRI 796 500 µm approximate posterior distribution.

### A.7 Posterior predictive checks from varying area

**Figure S17:**
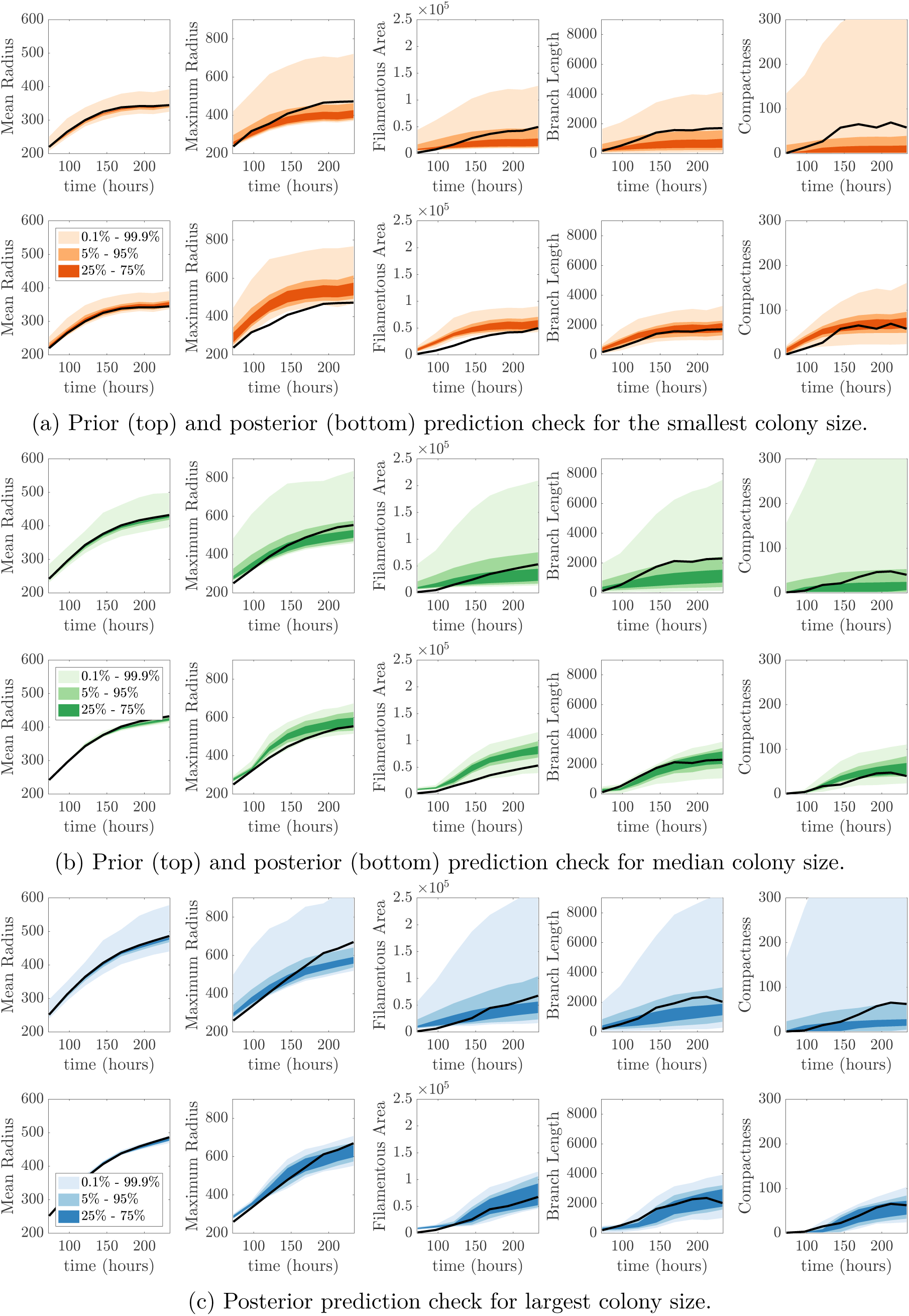
Prior (top) and posterior (bottom) prediction check for (a) smallest colony, (b) median colony and (c) largest colony based on area.

### A.8 Variability of area among replicates of the same experimental conditions

We present the colonies at the final experimental time to show the variability within an individual experiment of AWRI 796 50 µm as shown in Figure A.18.

**Figure S18:**
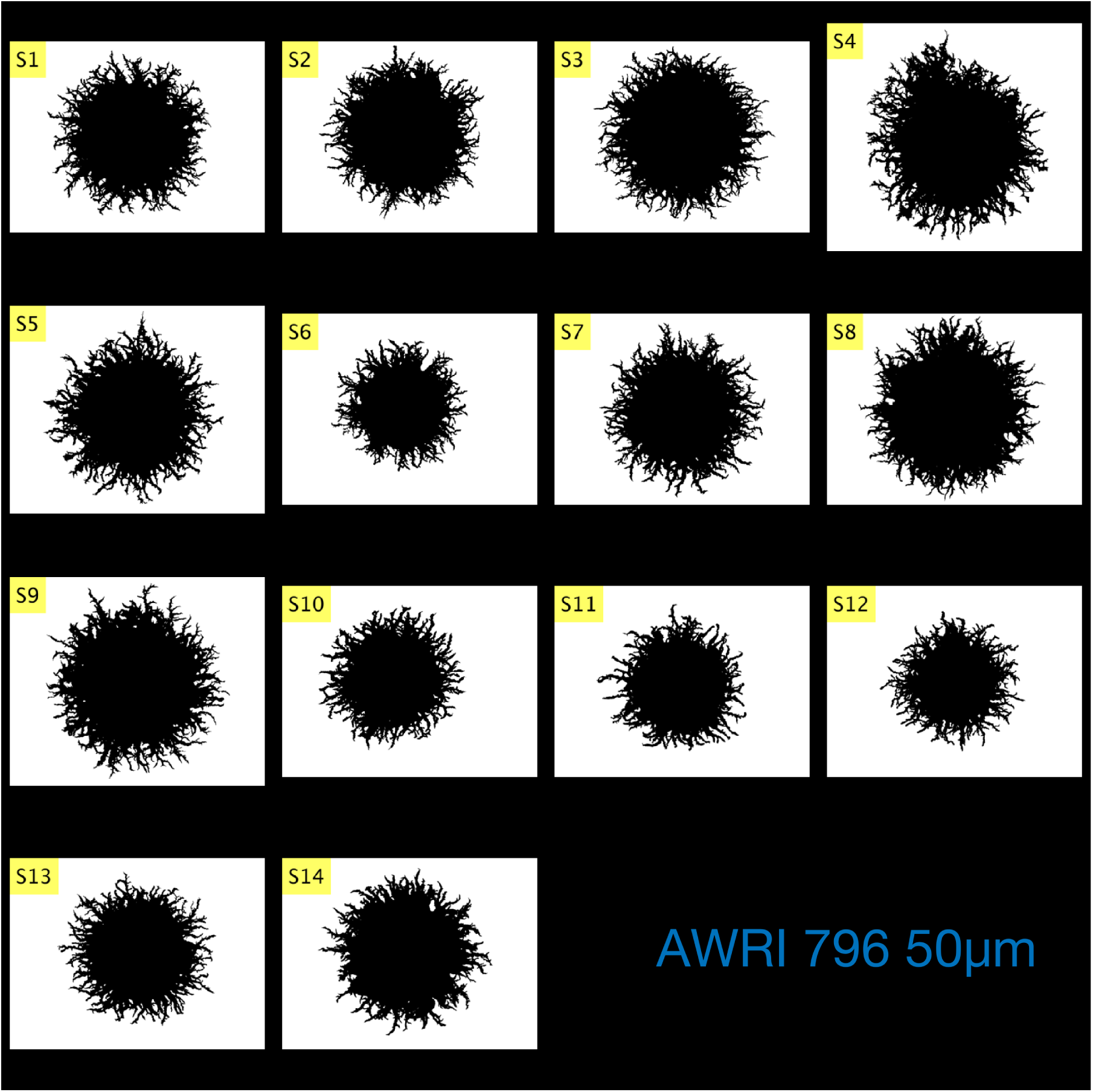
Variability of experimental image across the AWRI 796 50 µm experiments at final time (233 hours). The smallest colony was sample 12, the median size was sample 14, and the largest was sample 9.

